# Darwinian fitness, its directional derivative, and Hamilton’s rule for limited dispersal with class structure under within- and between-generation environmental stochasticity

**DOI:** 10.64898/2026.05.05.722983

**Authors:** Laurent Lehmann

**Affiliations:** Department of Ecology and Evolution, University of Lausanne, Switzerland

**Keywords:** Invasion fitness, stochastic growth rate, selection gradient, structured populations, stochastic fluctuations, relatedness, Hamilton’s rule, inclusive fitness theory

## Abstract

Darwinian fitness is equated here with invasion fitness and defined as that quantity determining the fate—certain extinction or possible spread—of a single mutant type. We derive it, together with its phenotypic derivative, for evolution occurring in group-structured populations under limited genetic mixing, where the demography of the focal species and its environment is modeled as a discrete-time stochastic process. Reproduction, physiological development, dispersal, and survival are influenced by interactions within and between groups and by environmental fluctuations within and across generations. Using multitype branching processes in random environments, we show that invasion fitness is predicted by a stochastic growth rate that can be represented biologically in two meaningful genealogical ways. First, as the long-term geometric mean of the expected per-capita number of mutant copies produced per time step by a representative member of the mutant lineage. Second, as the long-term geometric mean of the expected reproductive-value-weighted per-capita number of mutant copies produced by such an individual. This latter representation is useful for computing the phenotypic directional derivative of invasion fitness. Moreover, this derivative can be written as an actor-centered inclusive-fitness effect derived from properties of the resident population process alone. This effect depends on class-specific fitness differentials, relatedness, reproductive values, and class frequencies. However, unless generation- and class-specific fitness defines a stochastic matrix, the derivative does not separate stochastic reproductive values from relatedness and class frequencies, and must be evaluated by simulations. In summary, we formalize invasion fitness biologically quite generally and show how Hamilton’s marginal rule can be deduced from it.

## 1 Introduction

The selection gradient on a quantitative trait—the derivative of an appropriate measure of Darwinian fitness with respect to individual trait value—is a central quantity for understanding adaptation in evolutionary biology. By indicating the direction of trait change at any given population state, the selection gradient allows to ascertain directional selection and thus convergence stability under long-term evolution. From extraordinary sex ratios to the diversity of life histories and the evolution of social interactions and altruism, countless results about adaptation have been identified and understood through the use of phenotypic derivatives of fitness (for books and reviews illustrating the use of selection gradients to solve myriads of problems see, e.g. Charlesworth, 1994; Bulmer, 1994; Taylor and Frank, 1996; Frank, 1998; Geritz et al., 1998; Caswell, 2001; Rousset, 2004; Vincent and Brown, 2005; Otto and Day, 2007; Dercole and Rinaldi, 2008; Lehmann and Rousset, 2014; Van Cleve, 2015; Lion, 2018b; McNamara and Leimar, 2020; Avila and Mullon, 2023).

One aim of this paper is to derive the selection gradient for evolution occurring in a group-structured population and simultaneously accounting for all main heterogeneities individuals can be exposed to under the assumptions that the discrete-time and discrete state stochastic process underlying demographic, ecological and environmental fluctuations in the population is ergodic. In such a metapopulation, interacting individuals within and between groups may experience different group states and/or environmental conditions. The set of heterogeneities, or contexts, that an individual can experience can then be partitioned into between-generation heterogeneity and within-generation heterogeneity. Between-generation heterogeneity captures the different contexts triggered by global environmental variation and stochasticity across demographic time points, which affect all individuals in the population alike. Within-generation heterogeneity is more complex and encompasses a range of situations. These include stochasticity at the individual level and the physiological states in which individuals can reside (e.g., different ages, sexes, mating types, or stages). They also include the local biotic and abiotic conditions that affect all individuals within a group alike and vary across the population (e.g., group size, age structure, local resources, or number of predators). Finally, owing to the finite size of groups and limited genetic mixing, genetic fluctuations within groups imply that individuals are exposed to different numbers of other individuals who are identical-by-descent (IBD) to them. This invariably results in the occurrence of kin selection (e.g., Michod, 1982; Frank, 1998; Rousset, 2004) and in the kin selection literature heterogeneities structuring populations that are not the result of IBD assortative interactions are often referred to as class structure (e.g., Taylor, 1990; Frank, 1998; Rousset, 2004; Grafen, 2015; Priklopil and Lehmann, 2021).

The phenotypic selection gradient has been derived for each of the above types of situation separately, and in some work covering several situations together (e.g. Taylor, 1989; Tuljapurkar, 1990; Taylor and Frank, 1996; Frank, 1998; Ronce et al., 2000; Rousset, 2004; Rousset and Ronce, 2004; Lehmann and Balloux, 2007; Van Cleve, 2015; Lion, 2018a; Priklopil and Lehmann, 2021), yet never in a framework that simultaneously covers all heterogeneities and stochasticities. This leaves open the questions of what is the general biological representation of the phenotypic selection gradient in a group-structured population and how does within and between generation heterogeneity interact to affect directional kin selection.

Deriving selection gradients requires a measure of Darwinian fitness, which is here taken to be a measure determining whether a phenotypic mutation introduced as a single copy into a very large population—when everything else is held constant—will vanish with certainty or has a chance of persisting. This measure is here defined as invasion fitness and needs to account for all the different sources of heterogeneity and stochasticity. In populations subject to global environmental fluctuations but no within-generation stochasticity, invasion fitness can be taken to be the exponential of the stochastic growth rate of a mutant, which generalizes the geometric growth rate (or Malthusian parameter) to stochastic environments (e.g. Tuljapurkar, 1989, 1990; Metz et al., 1992; Rand et al., 1994; Arnold et al., 1994; Ferrière and Gatto, 1995). The selection gradient on a quantitative trait can then be obtained as the phenotypic derivative of this growth rate. From a mathematical perspective, this growth rate is obtained from a product of matrices when heterogeneities are discrete, as will be assumed here. This can make it difficult to (i) represent invasion fitness explicitly in terms of individual survival and reproductive components and (ii) calculate its derivative, especially in group-structured populations under limited genetic mixing, where the underlying state space becomes enormous owing to genetic fluctuations.

From an evolutionary perspective, one would thus like to have a biologically explicit representation of invasion fitness. Namely, an expression for the fitness of a mutant that depends explicitly on the individual-centered survival and reproductive components borne by mutant individuals. For instance, the Malthusian growth rate of a mutant depends on survival and reproduction, but this dependence is left implicit (since the Malthusian growth rate solves the Euler–Lotka equation, e.g. Charlesworth, 1994; Karlin and Taylor, 1981; Case, 2000). By contrast, the lifetime reproductive success of a mutant, which is a proxy for the Malthusian growth rate (Mode, 1968; Karlin and Taylor, 1981; Michod, 1979; Charlesworth, 1994; Case, 2000), is biologically explicit. It is the sum over all ages of the effective fertility at each age weighted by the probability of survival to that age. Having a biologically explicit representation of invasion fitness in terms of individual fitness components is desirable for gaining intuition and reasoning about the evolutionary process, and may also help in evaluating the selection gradient. Bulmer (1986) and Frank (1998) discuss how working directly with the raw and abstract growth rate has been problematic for understanding selection in the context of dispersal evolution.

The representation of invasion fitness in well-mixed class-structured populations was first formulated in terms of entropy (Demetrius, 1974, 1983; Arnold et al., 1994). In this framework, the growth rate is expressed as the sum of evolutionary entropy and a reproductive potential (Demetrius, 1983, eq. 4.19, Arnold et al., 1994, eq. 2.10). This representation is obtained by adopting a lineage (or genealogical) perspective, considering the reproduction and survival of the original mutant progenitor, as well as that of its descendants, the descendants of its descendants, and so on. In this way, invasion fitness is taken to be the lineage fitness of the mutant and the entropy representation is derived by considering events along the lineage backward in time. Specifically, one considers the probability that a randomly sampled lineage member descends from a mother in a particular class. These probabilities are then averaged over the difference between the reproductive potential of the parent of such a mother (involving the logarithm of class-specific fitness) and the degree of surprise (entropy) associated with producing such a mother (e.g., Demetrius, 1983, eq. 4.19). The lineage perspective can also be taken forward in time to yield another representation of invasion fitness. In this case, one considers the probability of sampling a lineage member in a given class and averages these probabilities over the corresponding class-specific fitness components. From this forward perspective, invasion fitness is simply the expected fitness of a randomly sampled lineage member (Lehmann et al., 2016, eq. 5, Van Cleve, 2023, eq. 6.3). This interpretation naturally incorporates within generation stochasticity and genetic fluctuations under limited mixing, since the fitness of an individual in a given class is itself an expected fitness averaged over all genetic backgrounds in which that individual may occur. Invasion fitness is thus inclusive, as it accounts for all effects arising from interactions among relatives. Invasion fitness can also be expressed as the reproductive-value-weighted average number of mutant copies produced per demographic time step by a randomly sampled lineage member (Lehmann et al., 2016, eq. C.5; Ohtsuki et al., 2020, eq. 16.a). This formulation is particularly convenient for deriving the various fitness derivatives used in invasion analyses of class-structured populations (Ohtsuki et al., 2020).

The goal of this paper is twofold. First, to derive invasion fitness and provide biological interpretations thereof under the forward-lineage perspective in metapopulations that incorporate all aforementioned heterogeneities. Second, to derive the directional derivative of invasion fitness when phenotypic traits can be expressed in a class-specific manner (i.e., phenotypic plasticity) and put it under the form of an actor-centered inclusive-fitness effect. To this end, the paper weaves together threads from several previous works (Tanny, 1981; Steinsaltz et al., 2011; Lehmann et al., 2016; Ohtsuki et al., 2020; Priklopil and Lehmann, 2021, 2024) and is organized as follows. It begins by outlining the model, including the underlying demographic process and population structure. Next, using classical results from multitype branching process theory, two biological representations of the invasion fitness of a mutant introduced as a single copy into a resident population are obtained. Finally, the directional derivative of invasion fitness is derived, bridging the standard expression for the derivative of the stochastic growth rate in fluctuating environments (Tuljapurkar, 1990; Steinsaltz et al., 2011) with Hamilton’s marginal rule for class-structured populations (Frank, 1998; Rousset, 2004; Priklopil and Lehmann, 2021).

## 2 Model

### 2.1 Biological setting

Consider a population of conspecific, asexually reproducing individuals inhabiting an infinite number of local groups. These groups are connected by random, uniform individual migration, i.e., the infinite island model of dispersal (Wright, 1931; Rousset, 2004). Each group may vary in size, but density-dependent regulation ensures that it contains a finite and discrete number of individuals (i.e., each group is bounded in size). The finite set of possible group sizes is denoted by *N* and includes the state with zero individuals (Tables 1– 3 summarize the main notation). Each individual within a group can occupy one of a finite number of physiological states, such as age and/or size. We denote by *P* the finite set of physiological states available to individuals. The demographic state of a group is defined as the collection of the physiological states of all individuals within that group and thus determines the number of individuals in each state. In a group of total size *N*, the demographic state is an element of *P*^*N*^, where *P*^*N*^ is the *N* th Cartesian product of *P*. Consequently, the demographic state space *D* of a group is given by *D* = ∪_*N*∈*N*_ *P*^*N*^, which is a finite set since group size is bounded.

**Table 1:** Definition of the different types of heterogeneities.

| Notations | Definitions |
| --- | --- |
| $\mathcal{P}$ | Finite set of physiological states an individual of the focal species can be in, e.g. age class, sex, stage. |
| $\mathcal{N}$ | Finite set $\{0, 1, 2, 3, 4, \dots, K\}$ of total number of individuals in a group, where $K$ is some finite maximum group size. |
| $\mathcal{R}$ | Finite set of local environmental conditions (abiotic or biotic) that individuals of the focal species can experience in a group (and that do not include demographic states of the focal species). |
| $\mathcal{E}$ | Finite set of global environmental conditions. |
| $\mathcal{D} = \cup_{N \in \mathcal{N}} \mathcal{P}^N$ | Set of local demographic states. |
| $\mathcal{S}_0 = \mathcal{R} \times \mathcal{D}$ | Set of local group states, which includes the empty group. |
| $\mathcal{S}$ | Set $\{s \in \mathcal{S}_0 \mid \exists a \in \mathcal{P} \text{ such that } N_{a\vert s} \geq 1\}$ of local group states excluding the empty group. |
| $\mathcal{C}$ | Set $\{(a, s) \in \mathcal{P} \times \mathcal{S} \mid N_{a\vert s} \geq 1\}$ of classes an individual can be in. The class of an individual is the state that determines its reproduction and survival in a given generation. The class space thus specifies the total within-generation heterogeneity. |
| $c = (a, s)$ | Element of the class state space $\mathcal{C}$, which consists of an individual in physiological state $a \in \mathcal{P}$ in a group in state $s \in \mathcal{S}$ such that $N_{a\vert s} \geq 1$. |
| $\mathcal{H} = \mathcal{E} \times \mathcal{C}$ | Set of different types of heterogeneities an individual can be exposed to, including within and between generation heterogeneity. |
| $N_{a\vert s}$ | Number of individuals in physiological state $a$ given a group is in state $s$. |
| $\mathcal{I}(s)$ | Set $\{\mathbf{i} \mid 0 \leq i_a \leq N_{a\vert s} \forall a \in \mathcal{P} \text{ and } \exists a \in \mathcal{P} \text{ such that } i_a \geq 1\}$ of all possible mutant copy number distributions in a group in state $s \in \mathcal{S}$ and such that there is at least a mutant present in the group. The vector $\mathbf{i} = (i_a)_{a \in \mathcal{P}}$ gives the number of mutant copies in each physiological state they can be in. |

**Table 2:** Definition of resident demographic quantities.

| Functions | Definitions |
| --- | --- |
| $p^\circ(e'\vert e)$ | Probability that a population in environmental state $e \in \mathcal{E}$ turns into state $e' \in \mathcal{E}$ in the next demographic time point. |
| $p^\circ(e)$ | Stationary probability that the population is in state $e \in \mathcal{E}$. |
| $p_b^\circ(e\vert e') = \frac{p^\circ(e'\vert e)p^\circ(e)}{p^\circ(e')}$ | Backward transition probability that given the population is taken in state $e' \in \mathcal{E}$ it was in state $e \in \mathcal{E}$ in the previous demographic time point. |
| $p^\circ(s', e' \vert s, e, \mathbf{p}_e^\circ)$ | Probability that a focal group in state $s$ in a population in state $e$ turns into a group in state $s'$ and the population into state $e'$ in the next demographic time point. |
| $p^\circ(s, e)$ | Stationary probability that a focal group experiences joint state $(s, e)$. |
| $\mathbf{p}_e^\circ = \{p^\circ(s \vert e)\}_{s \in \mathcal{S}_0}$ | Vector of conditional probabilities $p^\circ(s \vert e) = p^\circ(s, e)/p^\circ(e)$ that a group is in state $s$ given population state $e$ and that satisfy $\sum_{s \in \mathcal{S}_0} p^\circ(s \vert e) = 1$. |
| $w^\circ(c' \vert c, e)$ | Expected number of class $c'$ offspring produced by a class $c$ individual when the population is in state $e$. The argument $e$ therein entails that this is a function not only of $e$, but also of the distribution $\mathbf{p}_e^\circ$. |
| $z_{ase}$ | Resident quantitative trait of an individual in physiological state $a$ in a group in state $s$ in a population in state $e$. The profile of all possible traits is $\mathbf{z} = (z_h)_{h \in \mathcal{H}}$, which is a resident individual's reaction norm. |
| $w_{a's'\vert ase}(z_{\text{foc}}, \mathbf{z}_{\text{loc}}, \mathbf{z}_{\text{pop}})$ | Expected number of class $c' = (a', s')$ offspring produced by a focal individual in physiological class $a$ with trait $z_{\text{foc}}$ living in a group in state $s$ with trait profile $\mathbf{z}_{\text{loc}} = (z_{\text{loc}, a})_{a \in \mathcal{P}}$ in a population in state $e$ with trait profile $\mathbf{z}_{\text{pop}} = (z_{\text{pop}, c})_{c \in \mathcal{C}}$. Therein, $z_{\text{loc}, a}$ is the average trait among the focal's neighbors in state $a$ and $z_{\text{pop}, c}$ is the average population trait of class $c = (a, s)$ individuals. Expressed in terms of resident state specific traits the fitness function is $w_{a's'\vert ase}(z_{ase}, (z_{ase})_{a \in \mathcal{P}}, (z_{ce})_{c \in \mathcal{C}}) = w^\circ(a', s' \vert a, s, e)$. |
| $\bar{Z}_t^\circ(c)$ | Expected number of individuals of a resident lineage in class $c \in \mathcal{C}$ under a given realization of the environmental sequence experienced by the lineage. The total average number of individuals is $\bar{Z}_t^\circ = \sum_{c \in \mathcal{C}} \bar{Z}_t^\circ(c)$. |
| $Q_t^\circ(c)$ | Frequency of resident lineage members at time $t$ in class $c$ under a given realization of the environmental sequence experienced by the lineage. We have $Q_t^\circ(c) = \sum_{\mathbf{i} \in I(s)} Q_t^\circ(c, \mathbf{i})$, where $Q_t^\circ(c, \mathbf{i})$ is the frequency of lineage members at time $t$ in class $c$ and in a group with profile $\mathbf{i}$ of IBD residents. |
| $\tilde{V}_t^\circ(c)$ | Reproductive value of a resident individual of class $c$ at time $t$. |
| $V_t^\circ(c)$ | Normalized reproductive value $\tilde{V}_t^\circ(c) / (\sum_{c \in \mathcal{C}} \tilde{V}_t^\circ(c))$ of an individual of a resident lineage at time $t$ with the average normalized reproductive value being given by $\bar{V}_t^\circ = \sum_{c \in \mathcal{C}} V_t^\circ(c) Q_t^\circ(c)$. |
| $R_t^\circ(b \vert a, s)$ | Probability that, given a focal individual sampled in physiological state $a$ within a group in state $s$ at time $t$, another individual, randomly sampled in physiological state $b$ from the same group is identical-by-descent (IBD) to the focal. |
| $q^\circ(a, s \vert e)$ | Probability $\mathbb{E}[Q^\circ(a, s) \vert e]$ that a randomly sampled individual in a population in state $e$ is of class $(a, s)$. |
| $r^\circ(b \vert a, s, e)$ | Probability $\mathbb{E}[R^\circ(b \vert a, s) Q^\circ(a, s) \vert e] / q^\circ(a, s \vert e)$ that in a population in environmental state $e$ and given an individual has been sampled in class $(a, s)$, a randomly sampled neighbor of class $b$ is IBD to the former individual. |

**Table 3:** Definition of mutant lineage quantities.

| Functions | Definitions |
| --- | --- |
| $Z_t$ | Random number of individuals of the mutant lineage at time $t$. |
| $\bar{Z}_t(c)$ | Expected number of individuals of the mutant lineage in class $c \in \mathcal{C}$ at time $t$ under a given realization of the environmental sequence experienced by the lineage; namely $\bar{Z}_t(c) = \bar{Z}_t(a, s) = \sum_{\mathbf{i} \in I(s)} i_a \bar{Z}_t^g(s, \mathbf{i})$. The expected total lineage size is $\bar{Z}_t = \sum_{c \in \mathcal{C}} \bar{Z}_t(c)$. The expectation is thus taken over within generation stochasticity, but not between generation stochasticity. |
| $\mathbf{M}_t$ | Mean transition matrix of the mutant lineage with element $m(s', \mathbf{i}' \mid s, \mathbf{i}, e_t)$ giving the expected number of groups in local state $(s', \mathbf{i}') \in \{(s, \mathbf{i}) \mid s \in \mathcal{S} \text{ and } \mathbf{i} \in I(s)\}$ that are “produced” over one demographic time period at time $t$ by a focal group in state $(s, \mathbf{i}) \in \{(s, \mathbf{i}) \mid s \in \mathcal{S} \text{ and } \mathbf{i} \in I(s)\}$ when the population at large experiences environmental state $e_t \in \mathcal{E}$. |
| $Q_t(c, \mathbf{i})$ | Frequency of mutants at time $t$ in class $c$ and in a group with mutant profile $\mathbf{i}$. |
| $Q_t(c)$ | Frequency of mutants at time $t$ in class $c$; namely $Q_t(c) = \sum_{\mathbf{i} \in I(s)} Q_t(c, \mathbf{i})$. |
| $\bar{V}_t$ | Average (neutral) reproductive value $\sum_{c \in \mathcal{C}} V_t^\circ(c) Q_t(c)$ of a mutant lineage member at time $t$. |
| $\bar{Z}_t^g(s, \mathbf{i})$ | Mean number of groups in state $(s, \mathbf{i})$ at time $t$. |
| $\bar{\mathbf{z}}_t^g$ | Vector $(\bar{Z}_t^g(s, \mathbf{i}))_{s \in \mathcal{S}, \mathbf{i} \in I(s)}$ collecting the random variables $\bar{Z}_t^g(s, \mathbf{i})$. |
| $g = \log(W)$ | Growth rate of the mutant lineage. |
| $W$ | Invasion fitness of the mutant type: the mutant lineage introduced as a single copy into a very large population goes extinct with probability one if $W \leq 1$ and has a chance of persisting if $W > 1$, and in this latter case the asymptotic growth ratio of the mutant lineage is $W = \exp\left(\lim_{t \rightarrow \infty} \frac{1}{t} \log(Z_t)\right)$. |
| $w(a', s' \mid a, s, \mathbf{i}, e)$ | Expected number of successful offspring of class $a'$, which settle in groups in state $s'$, and are produced by a single mutant individual in class $a$ reproducing in a group in state $(s, \mathbf{i})$ in a population in global environmental state $e$ (eq. 15). |
| $\mathbf{z} + \delta \boldsymbol{\eta}$ | Mutant trait profile, where $\boldsymbol{\eta} = (\eta_h)_{h \in \mathcal{H}}$ is the mutant deviation profile and $\mathbf{z} = (z_h)_{h \in \mathcal{H}}$ is the resident trait profile. |

Demographic time is assumed to be discrete, meaning that group sizes and compositions are censused at regular time intervals. Between any two such time steps, the number of individuals in a group, their physiological states, and the overall group state may change due to individual-level stochastic processes such as reproduction, survival, development, and dispersal. Offspring number, adult survival, physiological state transitions, and dispersal are all modeled as random variables with finite means and variances. The sequence of these events may vary depending on the biological context and may be influenced by interactions among individuals within and between groups, as well as by local environmental conditions affecting all individuals within a group equally. To capture local environmental heterogeneity (e.g., resource availability, predator abundance), we denote by *ℛ* the finite set of local environmental conditions. The state space that a group can experience is then given by *S*_0_ = *ℛ* × *D*, the Cartesian product of the space of local environments *ℛ* and the demographic state space *D*. The state of a group determines the distribution of all biotic and abiotic features therein, including those of the focal species. The number of individuals in physiological state *a* ∈ *P* within a group in state *s* ∈ *S*_0_ is denoted by *N*_*a*|*s*_ and *S*_0_ includes the possibility that the group is extinct [i.e., the empty patch when *N*_*a*|*s*_ = 0 for all *a* ∈ *P*].

Finally, we assume that the demographic population process is also influenced by a global environment (e.g., annual temperature or precipitation), which is experienced uniformly by all individuals in the population and may change between demographic time steps. We denote by *ℰ* the finite set of global environmental states that population members can experience.

Let us now embed a population genetic process within this demographic population process, allowing different alleles to segregate at a given genetic locus. We assume that, in the absence of mutations, the entire demographic-environmental-genetic system converges to an attractor point, consistent with the external stability assumption of invasion analysis that we endorse (e.g. Eshel, 1996; Metz et al., 1992; Altenberg et al., 2017; Van Cleve, 2023). Now, suppose a mutant allele, coding for a new trait, arises within the resident population. Under what conditions will this mutant increase in frequency rather than go extinct? To answer this question, we need to characterize the resident population process. Therefore, we begin our analysis with that and assume without loss of generality that the resident population is monomorphic (allowing for resident polymorphism would only make the notation more cumbersome, but not affect any qualitative results, Priklopil and Lehmann, 2021).

### 2.2 The resident population

We refer to a monomorphic resident population (the resident population for short) as a population subject to the assumptions described in section 2.1 and where individuals are all genetically alike. The defining feature of a monomorphic resident population is that for each environmental state in *ℰ*, individual reproduction, survival, development, and dispersal are exchangeable random variables for all individuals experiencing the same physiological state in *P* and group state in *S*_0_. In a (monomorphic) resident population, the evolutionary process is thus a neutral process as conceived in population genetics (e.g., Crow and Kimura, 1970; Nagylaki, 1992; Ewens, 2004 and see in particular Cannings, 1975 for structured populations).

To formalize the resident population process, we can, by virtue of the island model assumptions in Section 2.1, where space is only implicitly considered, focus on the stochastic process that describes the change in state of a representative group, henceforth referred to as the focal group. The state changes of the focal group are assumed to be described by a non-homogeneous Markov chain that is irreducible and aperiodic on the state space *S*_0_. Let 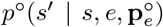 denote the non-homogeneous stationary transition probability of this chain. Specifically, this is the probability that the focal group transitions from state *s* ∈ *S*_0_ in a given time period (the “parental generation”) to state *s*^′^ ∈ *S*_0_ in the next demographic time period (the “offspring generation”), when the population is in environmental state *e* ∈ *E*. Throughout the analysis, the superscript ° indicates that the quantity is evaluated with trait characteristics of the resident population only. For simplicity, the traits of individuals in the population are not mentioned explicitly for now, as this is not required at this stage (traits will appear in section 4.1). The aforementioned non-homogeneity of the transition probability arises for two reasons.

- Coupling by dispersal, since groups are interconnected through individual migration. As a result, the demographic process within the focal group depends on the states of the population at large; namely, the population as a whole behaves as an infinite set of interacting Markov chains. Consequently, the transition probability 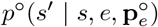 depends endogenously on the stationary distribution 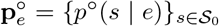, where *p*^°^(*s* | *e*) represents the conditional probability that a group resides in state *s*, given the population is in state *e*.
- Global environmental fluctuations, since the part of the environment that is experienced uniformly by all individuals in the population, changes between demographic time periods. This implies that the transition probability 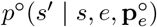 depends directly on *e*. The external process driving these environmental changes is modeled as a homogeneous, irreducible, and aperiodic Markov chain on the state space *E*, with transition probability *p*^°^(*e*^′^ | *e*) from state *e* to state *e*^′^. Thus, the stationary probability *p*^°^(*e*) *>* 0 that the population is in environmental state *e* ∈ *E* satisfies

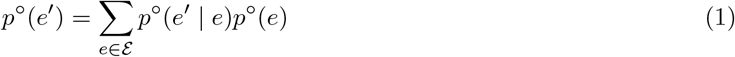

(e.g., Karlin and Taylor, 1975; Grimmett and Stirzaker, 2001).

In terms of *p*^°^(*e*^′^ | *e*) and 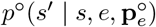, we can construct the transition probability

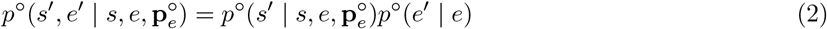

that the focal group in state *s* in a population in state *e* in the parental generation turns into a group in state *s*^′^ in a population in environmental state *e*^′^ in the offspring generation. Eq. (2) describes the dynamics of a bivariate non-homogeneous aperiodic irreducible Markov chain, which is an appropriate description of the environmental-demographic process, since the environmental process is itself time-homogeneous (Cogburn, 1980). The probability *p*^°^(*s*^′^, *e*^′^) that the focal group is in state *s*^′^ and the population is in state *e*^′^ satisfies

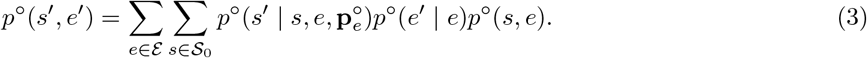

On knowledge of the stationary environmental process eq. (1), the conditional distribution 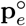 can be expressed in terms of the bivariate probabilities *p*^°^(*s, e*) since *p*^°^(*s* | *e*) = *p*^°^(*s, e*)*/p*^°^(*e*). Hence the bivariate distribution 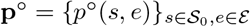 satisfies

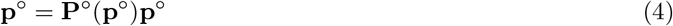

for a transition matrix **P**^°^(**p**^°^) with elements 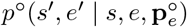 expressed in terms of elements of **p**^°^ by using 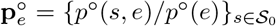 (which requires eq. 1). We assume that eq. (4) has a single stable equilibrium 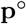 with a positive average group size, which in turn allows 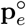 to be evaluated for all *e*.

Eq. (4) characterizes the distribution of group states in the (meta)-population process, accounting for local demographic and environmental stochasticity as well as global environmental stochasticity. For a single global environmental state, resident demographic dynamics, described by group transition probabilities that depend endogenously on the distribution of states in the population as a whole, have been used under numerous biological scenarios (e.g., Ronce et al., 2000; Metz and Gyllenberg, 2001; Cadet et al., 2003; Parvinen et al., 2003; Rousset and Ronce, 2004; Leturque and Rousset, 2004; Lehmann et al., 2006; Alizon and Taylor, 2008; Wild et al., 2009; Wild, 2011; Rodrigues and Gardner, 2012; Parvinen, 2013; Massol and Débarre, 2015; Rodrigues, 2018a; Kuijper and Johnstone, 2019; Priklopil and Lehmann, 2021). Models addressing global environmental change under limited dispersal have also been developed, though these typically consider groups of infinite size (Svardal et al., 2015) or operate within the framework of diffusion processes (Gillespie, 1991).

## 3 Invasion fitness: the raw and the cooked

### 3.1 Mutant lineage and resistance to invasion

We now investigate the invasion process of a mutant introduced as a single copy into the backdrop of the resident population at its demographic attractor characterized by the group state distribution eq. (4). To that end, we need two additional pieces of notation. First, let *I*(*s*) = {**i** | 0 ≤ *i*_*a*_ ≤ *N*_*a*|*s*_ ∀*a* ∈ *P* and ∃*a* ∈ *P* such that *i*_*a*_ ≥ 1} stand for the set of all possible mutant copy number distributions in a group in state *s* ∈ *S* with at least one mutant. Therein, the vector **i** = (*i*_*a*_)_*a*∈*P*_ gives the number of mutant copies in each physiological state they can be in. The number of mutant copies in physiological state *a* ∈ *P* in such a group state can thus range between 0 (all individuals are residents) and *N*_*a*|*s*_ (all individuals are mutants). Second, let *S* = {*s* ∈ *S*_0_ | ∃*a* ∈ *P* such that *N*_*a*|*s*_ ≥ 1} be the set of all group states excluding the extinct group (empty patch).

In order to describe the process of mutant copy number in the population, let *Z*_*t*_ denote the random number of individuals at time *t* descending from the single mutant ancestor (or progenitor) introduced at time *t* = 0 into the resident population. The mutant lineage size *Z*_*t*_ includes the progenitor when alive, its living direct offspring (including itself if it survives), the living offspring of its offspring, and so on through all subsequent generations. This number can be written as

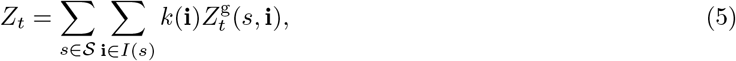

where

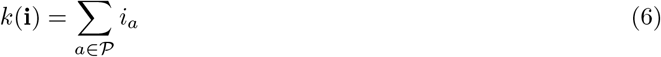

is the total number of mutant individuals in a group in genetic state **i** and 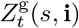 is the random number of groups in the population in state (*s*, **i**) and that contain at least one mutant.

The resident population will be said to be *resistant to invasion* by the mutant if the mutant lineage descending from the progenitor of the lineage goes extinct, i.e. *Z*_*t*_ = 0, with certainty for any physiological state *a* ∈ *P*, group state *s* ∈ *S*, and environmental state *e* ∈ *E*, in which the single progenitor of the mutant’s lineage can appear.

Because the total population size is infinite (i.e., the population consists of an infinite number of finite-sized groups), the dynamics of the mutant lineage is appropriately modeled as a branching process. The defining feature of a branching process is that a new, independent copy of the process begins each time a “new individual” is born (Harris, 1963; Mode, 1968, 1971; Kimmel and Axelrod, 2015) and we here consider a branching process in a random environment (e.g. Athreya and Karlin, 1971; Karlin and Taylor, 1981; Tanny, 1981). By “new individual” is meant a mutant colonizing a virgin group without mutants. This is conveniently modeled as a branching process at the level of the 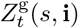 random variable, thus at the group level (e.g. Wild, 2011; Lehmann et al., 2016). By denoting 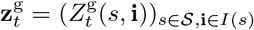 the vector collecting the 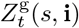 random variables, we see from eq. (5) that the event *Z*_*t*_ = 0 is equivalent to the event 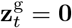.

To formalize resistance to invasion, let

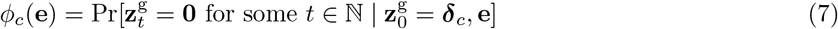

be the probability that the mutant lineage ultimately goes extinct, which is here conditioned on two events. First, the stationary environmental sequence to which the lineage is exposed is **e** = {*e*_0_, *e*_1_, *e*_2_, … }. This is induced by the transition probability *p*^°^(*e*_*t*+1_ | *e*_*t*_) and the distribution *p*^°^(*e*_0_) for the initial state *e*_0_ in which the progenitor appears (recall eq. 1). Second, the progenitor of the lineage appears in class *c* = (*a, s*) ∈ *C*, where the class of an individual combines its physiological state and the group state in which it resides and the class space is *C* = {(*a, s*) ∈ *P* × *S* | *N*_*a*|*s*_ ≥ 1}.^1^ When there are different classes of individuals at the same demographic time point in a population, individuals are divided into distinct categories (e.g., Cannings, 1975; Taylor, 1990; Frank, 1998; Rousset, 2004; Grafen, 2015; Lion, 2018a; Priklopil and Lehmann, 2021). The vector ***δ***_*c*_ has the same dimension as 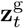 and contains one mutant individual in class *c* = (*a, s*) and zeros otherwise; that is, it represents a single mutant individual in physiological state *a* residing in a group in state *s*.

Since *ϕ*_*c*_(**e**) is a random variable in the range [0, 1] and owing to the fact that **e** is random, the probability of ultimate extinction of a lineage descending from a progenitor in class *c* is E[*ϕ*_*c*_(**e**)], where the expectation is over all possible environmental sequences that a lineage starting with a single copy can experience (e.g., Karlin and Taylor, 1975, p. 490). The resident population is thus resistant to invasion by the mutant if, for all *c* ∈ *C*, it holds that^2^

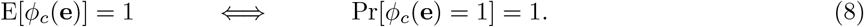

In order to characterize resistance to invasion, let us introduce the square matrix **M**_*t*_ whose (*s*^′^, **i**^′^; *s*, **i**) element, denoted *m*(*s*^′^, **i**^′^ | *s*, **i**, *e*_*t*_), is the expected number of groups in local state (*s*^′^, **i**^′^) that are “produced” over one demographic time period at time *t* by a focal group in state (*s*, **i**) when the population at large experiences environmental state *e*_*t*_ ∈ *E*. The dependence of *m*(*s*^′^, **i**^′^ | *s*, **i**, *e*_*t*_) on *e*_*t*_ is here a shorthand indicating that the expectation depends both directly on *e*_*t*_ and on the stationary group state distribution 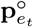 of the resident population at time *t* induced by *e*_*t*_, which affects any local group through dispersal (determined by eq. 3). This construction entails that the mutant does not affect the global environmental process, which can be affected only by the resident population.

We now endorse the following two standard assumptions about branching processes in random environments, which here describe mutant lineage dynamics. (i) There exists some time *t >* 0 such that any product **M**_*t*_ · · · **M**_0_ induced by its stationary environmental sequence **e** = {*e*_0_, *e*_1_, *e*_2_, …} has all entries positive and 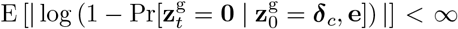 for some class *c*, where **0** is a vector of zeros (e.g. Tanny, 1981, p. 466). (ii) The mutant process is strongly regular (Tanny, 1981, Def. 9.1, p. 484). The branching process is not strongly regular if there exists a class *c* such that the process, when started with one mutant in that class, is sterile, and where the state entailing sterility may depend on the environmental sequence (Tanny, 1981, p. 483). Assumption (ii) is, therefore, a reasonable one from a biological perspective (yet menopause would be a counterexample). Assumption (i), meanwhile, implies an extinction probability dichotomy; namely, that the mutant branching process is such that Pr[*ϕ*_*c*_(**e**) = 1] = 1 or Pr[*ϕ*_*c*_(**e**) *<* 1] = 1 for all *c* ∈ *C* (Tanny, 1981, p. 466 and Th. 9.10, p. 488, Vatutin and Dyakonova, 2021, eq. 15). Biologically, this means that the mutant lineage either goes extinct with certainty or has a non-zero probability of invasion (i.e., any state with a finite number of mutants is transient). Finally, note that the first part of assumption (i), positivity of the matrix product along stationary ergodic environmental sequences, is the standard assumption of weak demographic ergodicity in evolutionary demography (e.g., Steinsaltz et al., 2011, p. 2).

It now follows from the classification results of multitype branching processes in random environments (Th. 9.6, p. 465 and Th. 9.10, p. 488 of Tanny, 1981 and see also Vatutin and Dyakonova, 2021, Th. 4) that if the multitype branching process describing the dynamics of the random variable 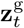 follows assumptions (i)-(ii), then the resident population is resistant to invasion by the mutant if and only if *g* ≤ 0, where

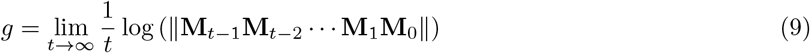

is a non-random growth rate and ∥·∥ is a matrix norm. Furthermore, conditional on non-certain extinction of the lineage, *g >* 0, one has 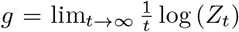 (Tanny, 1981, point (3) of Th. 9.6 along Def. 9.5 and Proposition 9.7).

The key feature of eq. (9) is that all matrices involved are mean matrices, whose entries represent expectations, despite the stochastic nature of individual reproduction, survival, maturation, and dispersal. Because *g* determines whether the mutant lineage goes extinct with certainty or has a chance of persisting in the population, the quantity *W* = exp(*g*) provides an appropriate measure of invasion fitness, i.e. the resident population is resistant to invasion by the mutant if and only if *W* ≤ 1. In the event of non-certain extinction, *g* is equivalent to the usual definition of the stochastic growth rate in evolutionary biology and demography (e.g. Tuljapurkar, 1989; Ferrière and Gatto, 1995; Metz et al., 1992), although the latter formalization typically ignores within-generation stochasticity. The interpretation of *W* is as the long-term geometric mean of the expected per-capita number of mutant copies produced per demographic time step by a randomly sampled lineage member. We now proceed with a series of rearrangements to make this biological interpretation explicit.

### 3.2 Invasion fitness in terms of average individual fitness

It follows from Tuljapurkar (1990, eq. 4.1.1 and eq. 4.2.2) that eq. (9) can be written as

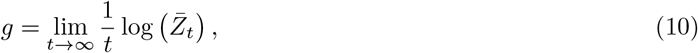

with

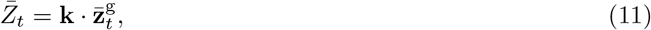

where · stands for the dot product between two vectors, the vector **k** = {*k*(*s*, **i**)}_*s*∈*S*,**i**∈*I*(*s*)_ has entries

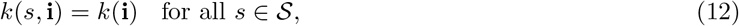

and the vector 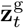 satisfies the recursion

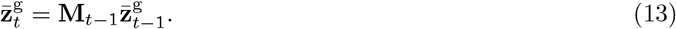

Here, 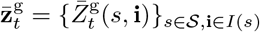 has element 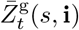, which gives the mean number of groups inhabited by the mutant lineage and in state (*s*, **i**) at time *t*. This is a random variable since the sequence of mean matrices {**M**_0_, **M**_1_, **M**_2_, ..} is random and depends on the realized environmental sequence individuals from the mutant lineage experience. Eq. (11) shows that 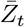 is the expected number of mutant copies at time *t* under this environmental sequence.

This expected number can be expressed as

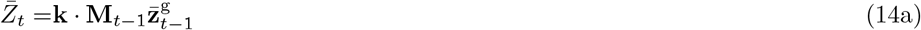

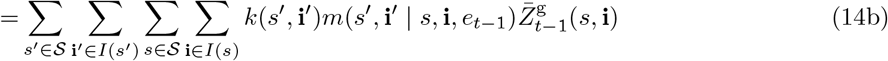

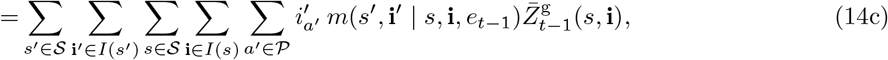

where eq. (6) and eq. (12) were used to obtain the last equality. Now observe that for all *s* ∈ *S*, **i** ∈ *I*(*s*), and *e* ∈ *E*, we have

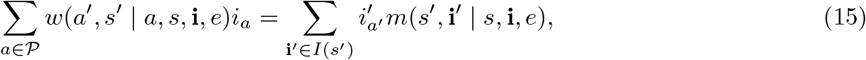

where *w*(*a*^′^, *s*^′^ | *a, s*, **i**, *e*) is the expected number of successful offspring of class *a*^′^ that settle in groups in local state *s*^′^ and are produced by a single mutant individual in state *a* residing in a group in state (*s*, **i**) in a population in global environmental state *e*. As is the case for *m*(*s*^′^, **i**^′^ | *s*, **i**, *e*), this dependence on *e* means that *w*(*a*^′^, *s*^′^ | *a, s*, **i**, *e*) depends both directly on *e* and on the stationary group state distribution 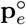. Eq. (15) holds since its right hand side is the total expected number of mutant individuals in a group of type *s*^′^ produced by a group of type (*s*, **i**) and generalizes previous relationships between individual fitness and group-state transitions to global environmental fluctuations (Lehmann et al., 2016, eq. A.5 and eq. F.1).

Inserting eq. (15) into the last line of eq. (14) and noting that *w*(*a*^′^, *s*^′^ | *a, s*, **i**, *e*) is zero unless *N*_*a*|*s*_ ≥ 1 and 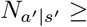 1 allows us to replace double sums of the form Σ_*a*∈*P*_ Σ_*s*∈*S*_ therein by Σ_(*a,s*)∈*C*_. This produces

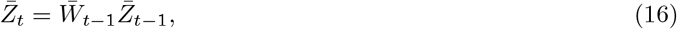

where

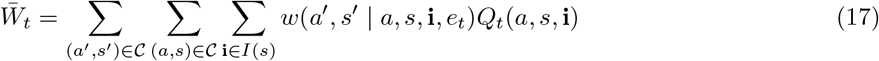

and

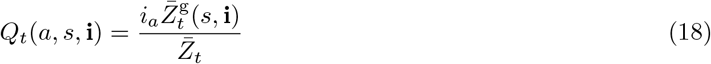

is the probability that a mutant randomly sampled from the average number of mutants at time *t* finds itself in physiological state *a* and in group state (*s*, **i**). Namely, we have Σ_(*a,s*)∈*C*_ Σ_**i**∈*I*(*s*)_ *Q*_*t*_(*a, s*, **i**) = 1 for all *t*, since 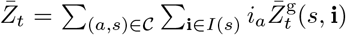. Hence, eq. (17) shows that 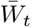 is the average of the expected fitness of a mutant at time *t*, averaged over all conditions a mutant copy can reside in; namely, 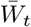 is the expected fitness of a randomly sampled mutant from the mutant lineage at time *t*.

Iterating eq. (16) given initial condition 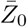 produces

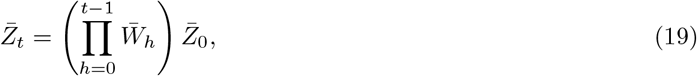

and substituting into eq. (10), we get 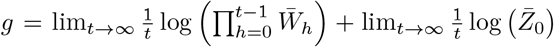. Since the initial condition is 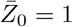, we have 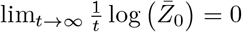 and can conclude that

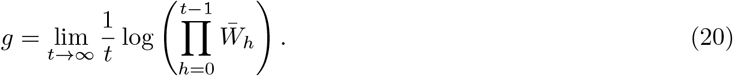

The exponential *W* = exp(*g*) then gives a representation of invasion fitness of the mutant as the geometric growth ratio:

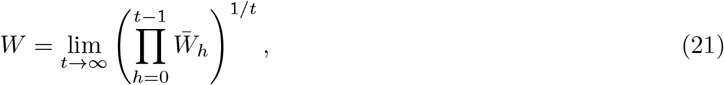

which holds since the exponential function is continuous and the stochastic growth rate is of the form *g* = lim_*t*→∞_ *x*_*t*_ whereby *W* = exp(lim_*t*→∞_ *x*_*t*_) = lim_*t*→∞_ exp(*x*_*t*_) (e.g. Apostol, 1974, Theorem 4.16). Invasion fitness *W* is thus the grand mean—a mean taken geometrically over between generation stochasticity and arithmetically across within generation heterogeneity—of the expected number of individuals produced by a single representative mutant lineage member.

We now add one additional step to the calculations. Let **Q**_*t*_ = {*Q*_*t*_(*c*, **i**)}_*c*∈*C*,**i**∈*I*(*s*)_ stand for the vector describing the demographic-genetic structure of a group. Owing to weak demographic ergodicity, the bivariate random variable (**Q**_*t*_, *e*_*t*_) is Markovian and will converge in distribution as *t* → ∞ and has well defined distribution and expectations (e.g. Steinsaltz et al., 2011)^3^. In this asymptotic distribution, there is a conditional distribution of the random variable **Q** given that environment *e* was obtained (e.g. Steinsaltz et al., 2011, p. 4). Hence, applying the ergodic theorem (e.g., Karlin and Taylor, 1975, Theorem 5.6, p. 487) to eq. (20), the stochastic growth rate has the representation

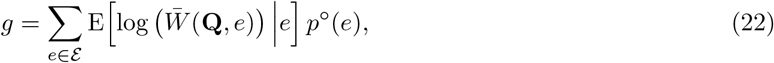

where E[·|*e*] is the conditional expectation over the distribution of **Q** given *e*, and

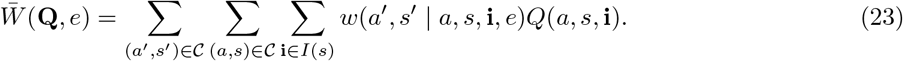

Thus *W* can be interpreted as the geometric mean over the joint environmental and demographic-genetic state distribution of the average expected per-capita number of successful offspring produced by a randomly sampled mutant lineage member. It thus describes the geometric mean of the expected individual fitness of a representative member of the mutant lineage, when the mutant reproductive process has reached stationarity in the resident population.

In the absence of class and genetic structure, i.e. *w*(*a*^′^, *s*^′^ | *a, s*, **i**, *e*) = *w*(*e*), eqs. (22)–(23) reduce to eq. (6) of Haccou and Iwasa (1995). In the absence of physiological structure and environmental fluctuations but with group and genetic structure, i.e. *w*(*a*^′^, *s*^′^ | *a, s*, **i**, *e*) = *w*(*s*^′^ | *s*, **i**), eqs. (22)–(23) reduce to eq. (5) of Lehmann et al. (2016). In the absence of group and genetic structure, and further without within generation stochasticity, but allowing for environmental fluctuations and physiological structure, Metz (2014, p. 7) provides an expression analogous to eq. (22) for a continuous-time process [while Arnold et al. (1994, Th. 4.3) provide a corresponding representation in terms of entropy for a discrete-time process]. Eqs. (22)–(23) thus extend these previous representations by simultaneously incorporating within and between generation stochasticity, physiological structure, and group and genetic structure within a single forward-lineage formulation of invasion fitness. We next provide a second representation of invasion fitness.

### 3.3 Invasion fitness in terms of reproductive value weighted fitness

Let us return to a monomorphic resident population where the evolutionary process is neutral, meaning that individuals within the same class are all exchangeable (Cannings, 1975). For this case, individual fitness *w*^°^(*c*^′^ | *c, e*) giving the expected number of successful offspring of class *c*^′^ = (*a*^′^, *s*^′^) produced by a single individual in class *c* = (*a, s*) in a population in global environmental state satisfies

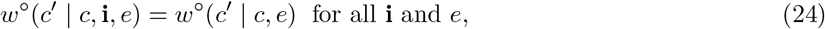

since the class specific fitness of all individuals within a given class must be the same and independent of the mutant distribution **i** (see Box 1 for an example of such fitness). This neutral individual fitness allows to introduce the neutral reproductive value 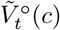 of an individual of class *c* of a resident lineage, defined to satisfy the recursion

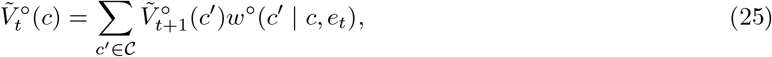

which in turn allows to define the normalized neutral reproductive value

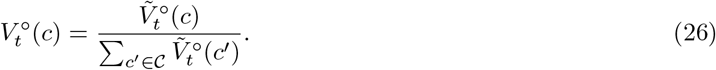

Unnormalized reproductive values are typically used in the analysis of natural selection in the absence of between-generation fluctuations (e.g. Taylor, 1990; Taylor and Frank, 1996; Rousset, 2004; Grafen, 2006; Priklopil and Lehmann, 2024), whereas a distinguishing feature of normalized reproductive values, which are typically used in the presence of between-generation fluctuations (e.g. Tuljapurkar, 1989, 1990; Lion, 2018a), is that, under the assumption of weak demographic stochasticity, they converge in distribution (e.g. Hennion, 1997; Steinsaltz et al., 2011).

Using the normalized reproductive value, let

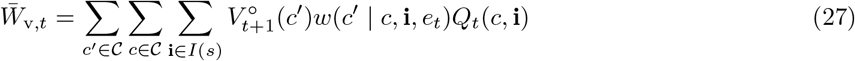

denote the reproductive-value-weighted mutant average offspring number at time *t*. Finally, let

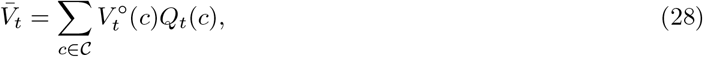

stand for the average (normalized) reproductive value of a mutant at time *t*, where *Q*_*t*_(*c*) = Σ_**i**∈*I*(*s*)_ *Q*_*t*_(*c*, **i**).

We now generalize a relationship between 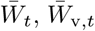, and 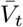 previously derived under panmixia for unnormalized reproductive values (Priklopil and Lehmann, 2024, eq. S.48). This is obtained by substituting eq. (18) into eq. (27) to yield

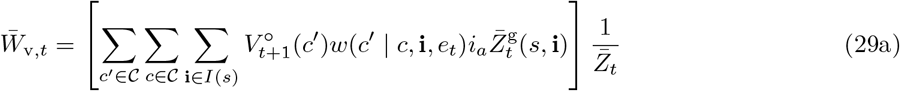

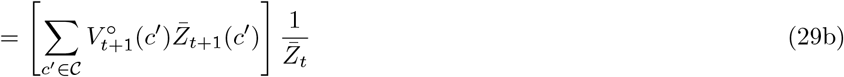

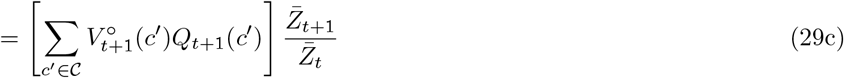

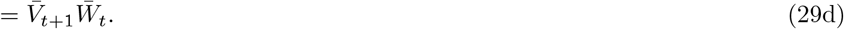

The second equality follows noting that 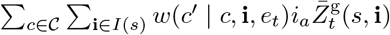 is the average number 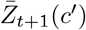 of mutants in class *c*^′^ at time *t* + 1, the third since 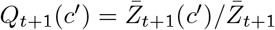, and the last equality by using eq. (28).

Using eq. (29d), the product appearing in the growth rate eq. (20) is

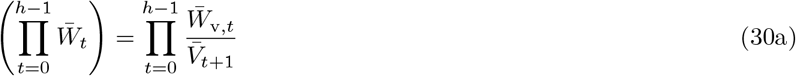

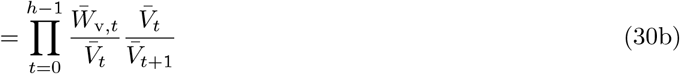

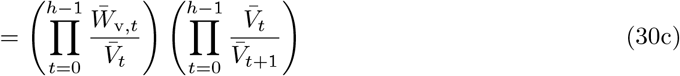

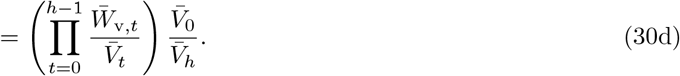

Substituting the last line into eq. (20) gives 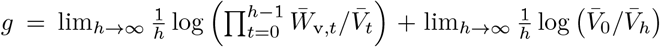. Because reproductive values are normalized according to eq. (26), a finite strictly positive terminal condition can be chosen for the backward recursion (25). In particular, one may set 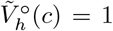 for all *c* ∈ *C*. This choice is arbitrary and does not affect the asymptotic normalized reproductive-value process since under weak demographic ergodicity 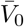 converges as the terminal horizon *h* is moved into the future. Hence, log 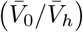 is a bounded term that does not grow linearly with the terminal horizon and 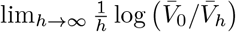 vanishes. Thus,

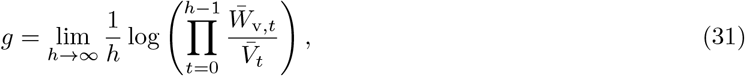

which in turn gives the invasion fitness representation as

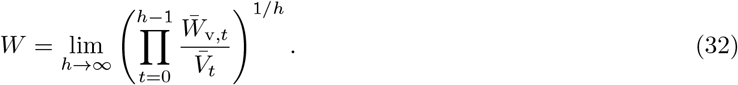

The per-generation term, 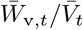, is the average reproductive value of the offspring of an individual randomly sampled from the mutant lineage at time *t*, relative to the reproductive value of that individual. Eq. (32) can also be written in a form analogous to eq. (23) by invoking the ergodic theorem and the fact that the normalized reproductive values converge in distribution, but the expectation is now taken over the neutral stochastic reproductive values and class frequencies. Invasion fitness thus also describes the geometric mean of the ratio between the average expected offspring reproductive value and the average expected reproductive value of a representative member of the mutant lineage, where both averages are taken over physiological, genetic, and group states, once the mutant reproductive process has reached stationarity in the resident population.

In the absence of physiological structure and environmental fluctuations, i.e. *w*(*a*^′^, *s*^′^ | *a, s*, **i**, *e*) = *w*(*s*^′^ | *s*, **i**), eq. (32) reduces to eq. (C.5) of Lehmann et al. (2016), but with the difference that unnormalized reproductive values were used. Likewise, in the absence of group and genetic structure, Priklopil and Lehmann (2024) provide an analogous finite-time representation of the effect of natural selection as a geometric mean of reproductive-value-weighted individual fitness, where the ratio 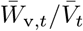 is expressed in terms of unnormalized reproductive values and gives precisely the measure of fitness that determines the change in mutant number due solely to natural selection during demographic time step *t* (Priklopil and Lehmann, 2024, eq. 9). This feature is lost in eq. (32). What is gained, however, is convergence in distribution of the normalized reproductive values, with *W* determining whether the lineage of a mutant introduced as a single copy vanishes with certainty or has a chance of increasing in number; conditional on such an increase, its asymptotic growth rate is *g* = log(*W*).

## 4 Derivative of invasion fitness

### 4.1 Derivative of individual fitness

We now evaluate the derivative d*g/* d*δ* at *δ* = 0. This is to be read as the infinitesimal effect of a mutant individual expressing trait deviation vector ***η*** = (*η*_*h*_)_*h*∈*H*_ when individuals in the resident population express trait **z** = (*z*_*h*_)_*h*∈*H*_, whereby the trait of a mutant is **z** + *δ****η***. Here, *H* denotes the set of all heterogeneities or contexts an individual can experience, partitioned as the Cartesian product *H* = *E* ×*C* of between-generation heterogeneity *E* and within-generation heterogeneity *C* experienced by an individual. It is well established that trait expression is often context-dependent, with a specific quantitative trait being possibly associated with each context an individual can experience (West-Eberhard, 1989). This is here captured by the fact that the trait a resident individual expressed in context *h* ∈ *H* is *z*_*h*_ ∈ ℝ, while the trait of the mutant is *z*_*h*_ + *δη*_*h*_ ∈ ℝ.

Given the focus on phenotypic effects, let us make explicit that individual fitness *w* should be conceptualized as being determined by the traits of interacting individuals. In a resident population with resident profile **z**, we now assume that individual fitness is of the form

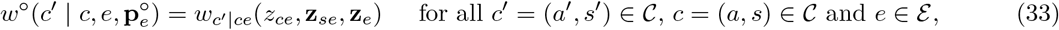

where **z**_*se*_ = (*z*_*ase*_)_*a*∈*P*_ and **z**_*e*_ = (*z*_*ce*_)_*c*∈*C*_. Here,

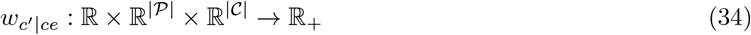

is an individual fitness function such that 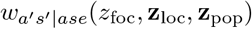 is the expected number of class (*a*^′^, *s*^′^) offspring produced by a focal individual of class (*a, s*) in a population in state *e*, where this individual has trait *z*_foc_; a group neighbour in state *a* has trait *z*_loc,*a*_, whereby the trait neighbour profile is **z**_loc_ = (*z*_loc,*a*_)_*a*∈*P*_ ∈ ℝ^|*P*|^; while an average population member of class *c* has trait *z*_pop,*c*_, whereby the trait population profile is **z**_pop_ = (*z*_pop,*c*_)_*c*∈*C*_ ∈ ℝ^|*C*|^. All individuals in the resident population in the same state express the same resident trait. Thus (*z*_loc,*a*_)_*a*∈*P*_ = (*z*_*ase*_)_*a*∈*P*_ and (*z*_pop,*c*_)_*c*∈*C*_ = (*z*_*ce*_)_*c*∈*C*_, but the trait notation in the fitness function 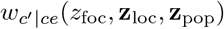 is useful as labels to distinguish the spatial relationship of individuals to the focal individual in the resident population and to compute fitness derivatives out of neutrality when individuals express different traits. Recall also that fitness 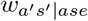, through the argument *e*, depends on the whole distribution **p**^°^ characterizing the resident demography. This in turn is likely to depend on the whole profile of traits (*z*_*h*_)_*h*∈*H*_ ∈ ℝ^|*H*|^, but we leave this dependence implicit as it will not be needed.

Equation (34) is an instantiation of the concept of an individual fitness function, which is widely used in the direct fitness method of kin selection theory (e.g. Taylor and Frank, 1996; Frank, 1998; Rousset, 2004; Rousset and Ronce, 2004; El Mouden and Gardner, 2008; Gardner, 2010; Ronce and Promislow, 2010; Koykka and Wild, 2015; Van Cleve, 2015; Rodrigues, 2018b; Kuijper and Johnstone, 2019; Avila and Mullon, 2023). Eq. (33) captures the three nested levels of heterogeneities individuals can experience in a metapopulation: from global population variation (*E*) to local group variation (*S*) and individual specific variation (*P*). Examples of individual fitness functions when *E* is singleton are very abundant in the literature. For instance, there are examples where (i) *P* is singleton and the group state space corresponds to different demographic group sizes, whereby *S* = *N* (e.g. Rousset and Ronce, 2004, eqs. 31-32, Lehmann et al., 2006, eqs. A8-A9), (ii) *P* is singleton and the group state space corresponds to different group level heterogeneities within generations, whereby *S* = *ℛ* (e.g., Rodrigues and Gardner, 2012, eqs. A4-A6, Kuijper and Johnstone, 2019, eq. 2.2, Ohtsuki et al., 2020, eqs. 45-47 and see eq. 60 for an example of this vein), (iii) *P* is non-singleton yet *S* is singleton (Ronce and Promislow, 2010, eqs. 3-4), (iv) neither *P* nor *S* are singleton, whereby the group state space *S* = *D* corresponds to a combination of physiological and demographic structure (e.g., Ronce et al., 2000, eqs. 3-4).

Let us now turn to mutant individual fitness *w*(*c*^′^ | *c*, **i**, *e*), which is understood as the fitness of a mutant with trait *z*_*ase*_ + *δη*_*ase*_ in a group in state (**i**, *s*) where the group distribution of traits is induced by that state. This means that each mutant individual in physiological state *a* in that group has trait *z*_*ase*_ + *δη*_*ase*_, while each resident individual has *z*_*ase*_ (how many of each is determined by **i**), and we assume that *w*(*c*^′^ | *c*, **i**, *e*) depends only on average traits (see Rousset, 2004, pp. 95-96 for why when computing first-order derivatives, considering only average phenotypes is sufficient in an evolutionary analysis). We will only need the derivative d*w*(*c*^′^ | *c*, **i**, *e*)*/* d*δ* = d*w*(*a*^′^, *s*^′^ | *a, s*, **i**, *e*)*/* d*δ* (evaluated at *δ* = 0). This can then be expressed in terms of eq. (34) as

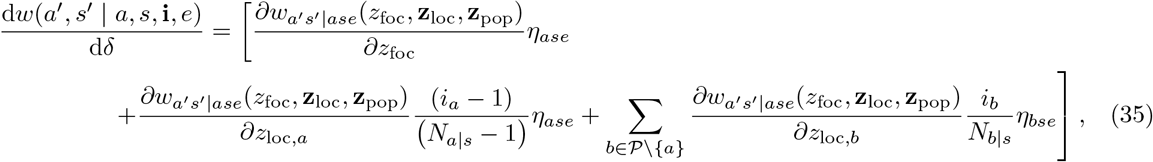

where all derivatives are evaluated at the resident trait profile (*z*_*ase*_)_*a*∈*P,s*∈*S*_ (since the total derivative is evaluated at *δ* = 0) and any sum over *b* ∈ *P* associated with a given group state *s* is from now on understood to run only over physiological states such that *N*_*b*|*s*_ ≥ 1, equivalently (*b, s*) ∈ *C* (and the second term on the right hand side is zero if *N*_*a*|*s*_ = 1). Eq. (35) holds because all individuals within the same class carrying the mutant allele are exchangeable. Thus, each mutant in class *b* expressing the mutant allele, expresses a phenotypic deviation *η*_*bse*_, which alters the fitness of any recipient of class *a* in the same way, indicating by a corresponding partial derivative evaluated in the resident population in eq. (35). And the effect of each such mutant is relative to the number of actors on a given recipient, since fitness depends only on average phenotypes. Adding up the effect of each actor on the average phenotype then gives eq. (35).

### 4.2 Directional derivative

We now have all the elements needed to evaluate the derivative d*g/* d*δ* of the stochastic growth rate; the analyticity of this derivative was established in Ruelle (1979). From eq. (31), this derivative can be computed as

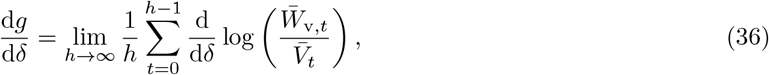

where remember that here and throughout all derivatives are evaluated at *δ* = 0, thus under neutrality; namely, at “°”. Eq. (36) assumes that differentiation and summation can be interchanged, which is valid if the right hand side of eq. (36) converges uniformly, as guaranteed by Apostol (1974, Theorem 9.14). Since invasion fitness is *W* = exp(*g*), whereby 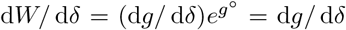 when evaluated at *δ* = 0, eq. (36) and the forthcoming result can also be read as the derivative of invasion fitness (eq. 21 or eq. 32).

Writing

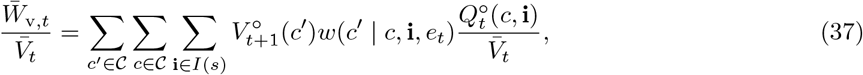

and using eq. (24) along the chain rule yields

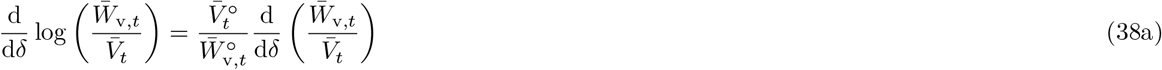

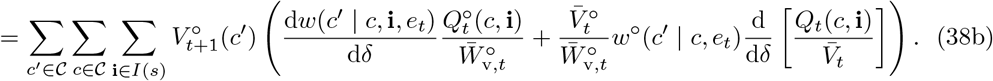

Now observe that

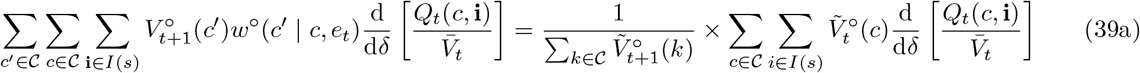

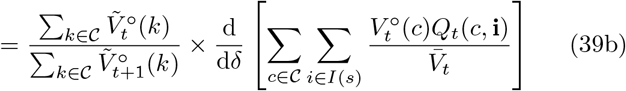

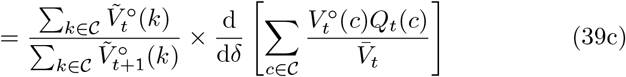

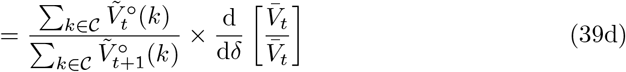

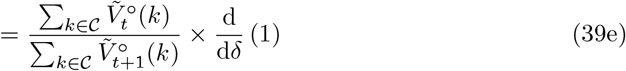

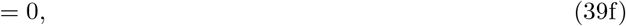

where eq. (26) was used to denormalize the reproductive value and then eq. (25) to obtain the first equality, eq. (26) was used to renormalize the reproductive value in the second equality, Σ _*i*∈*I*(*s*)_ *Q*_*t*_(*c*, **i**) = *Q*_*t*_(*c*) in the third equality, and eq. (28) in the fourth equality. Thus, eq. (38) reduces to

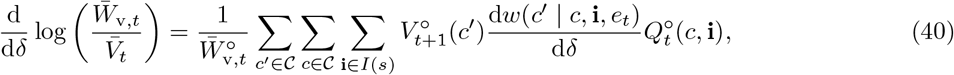

and substituting into eq. (36) produces

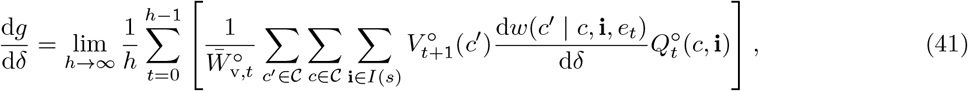

which, on insertion of eq. (35), finally gives

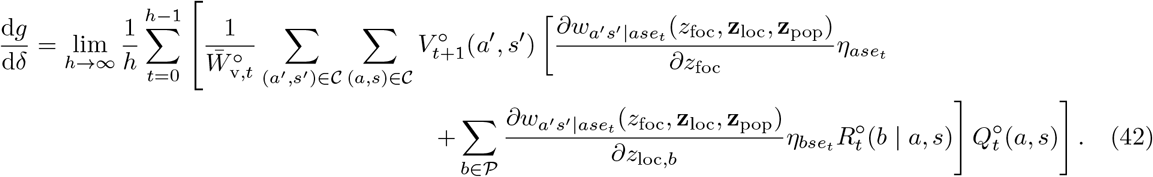

Here,

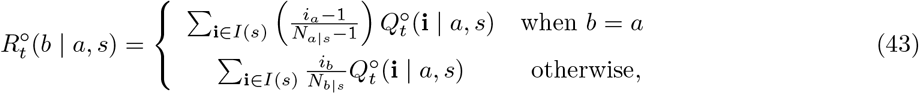

where

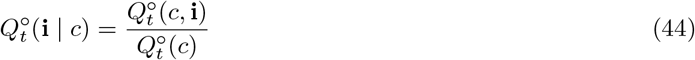

is the probability of genetic configuration **i** given that an individual has been sampled in class *c* = (*a, s*) [and with the convention that the *b* = *a* term in eq. (43) is zero when *N*_*a*|*s*_ = 1). Accordingly, 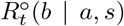 is the probability that, in a resident population, and given a focal individual is sampled in physiological state *a* within a group in state *s* at time *t*, another individual, randomly sampled in physiological state *b* from the same group, carries the same allele (and thus trait) IBD to the focal individual, i.e. both individuals ultimately descend from the progenitor of the mutant lineage (and this relatedness is defined only for *b* such that (*b, s*) ∈ *C*).

In eq. (42), the reproductive values do not depend on the distribution of mutant copy number: they are fully determined by the resident population and can be obtained from recursion equations that are of much lower dimension than the state space of the original branching process (e.g. eq. 25). In Appendix A, we show that this also holds for class frequencies, which satisfy

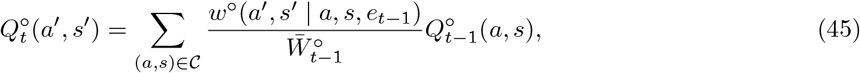

where 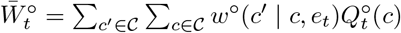, and for the relatedness coefficients, which satisfy

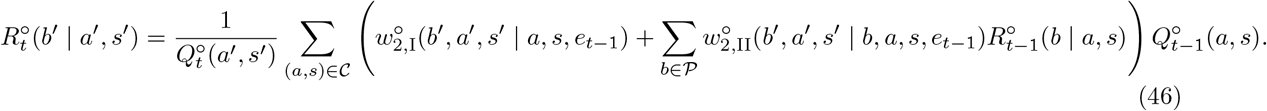

Here, 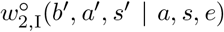 is the expected number of offspring in class (*a*^′^, *s*^′^) that descend from a parent reproducing in class (*a, s*) and that have a random neighbor in physiological state *b*^′^ that also descends from that parent (where all this is defined in the neutral resident process). Namely, this fitness measures the number of sibling pairs in physiological states *a*^′^ and *b*^′^ produced by a focal parent, which is an example of a generalized fitness function introduced in Ohtsuki et al. (2020). Meanwhile, fitness component 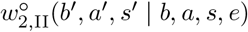 is the expected number of offspring in class (*a*^′^, *s*^′^) that descend from a focal parent in class (*a, s*) and that have a randomly sampled neighbor in physiological state *b*^′^, itself descending from any neighbor, in physiological state *b*, of the focal parent (see eq. A-10 for further explanation).

Suppose there are no interactions between relatives so that 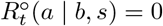 for all *a, b* and *s*. Additionally, set all phenotypic deviations to *η*_*ase*_ = 1. This scenario corresponds to evolution in a panmictic population where individuals across all conditions express the same trait value, *z*_foc_ = *z*_foc,*ase*_ (i.e., no phenotypic plasticity), a common assumption in the literature of adaptive dynamics in class-structured populations (e.g., Rousset, 2004, chapter 8). Then, eq. (42) is similar to the second line of eq. (11.2.7) of Tuljapurkar (1990) but here the biological model incorporates within-generation stochasticity. Eq. (42) thus extends the derivative of the stochastic growth rate in fluctuating environments to populations with limited genetic mixing and interactions between relatives.

### 4.3 Directional derivative as actor-centered inclusive fitness effect

Let 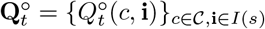 and 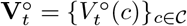 stand for the vectors collecting the random variables over all classes (and genetic states for 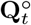). The bivariate random variables 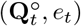 and 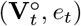 are Markovian random variables (i.e., they satisfy the Markov property), respectively, forward and backward in time, since the stationary environmental process **e** = {*e*_0_, *e*_1_, *e*_2_, …} is Markovian and the one step change in 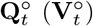 depends only on 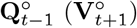 and is independent of 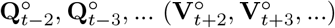. Both bivariate random variables will converge as *t* → ∞ and since the process is Markovian, if 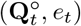 and 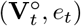 are chosen, respectively, from their (stationary) bivariate marginal distributions, then 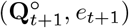 and 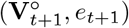 will also be chosen from the same marginal distributions. Accordingly, for each environmental state *e* that was picked in the stationary regime there exist conditional distributions for **Q**^°^ and **V**^°^, which are independent given *e*. All this is taken from Steinsaltz et al. (2011, p. 4), which applies to the present setting since the environmental-demographic-genetic state space is finite.

Applying the ergodic theorem (e.g., Karlin and Taylor, 1975, Theorem 5.6, p. 487) to eq. (42) and using the above convergence argument, we get

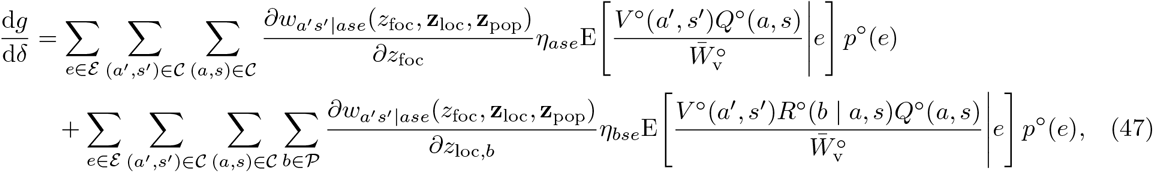

where

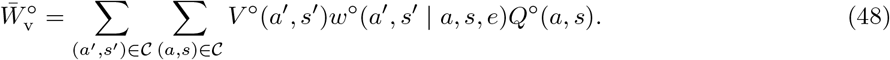

Since state *s* determines the number *N*_*a*|*s*_ of individuals in each class, the following relationship must hold:

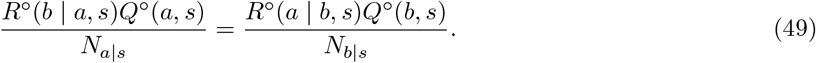

This can be understood by considering the probability of the event that in a population in state *e* with realized class frequencies {*Q*^°^(*c*)}_*c*∈*C*_ and relatedness profile {*R*^°^(*b* | *c*)}_*b*∈*P,c*∈*C*_, two randomly sampled gene copies, one in a state *a* individual the other in a state *b* individual and both in the same group in state *s*, are IBD. With probability *Q*^°^(*a, s*), the first sampled gene copy is an individual in class (*a, s*). In this case, the probability that the second gene comes from an individual in physiological state *b*, is *N*_*b*|*s*_*/*Σ_*a*∈P_ *N*_*a*|*s*_. Then, a randomly sampled *b* individual is IBD with the former individual with probability *R*^°^(*b* | *a, s*), whereby the probability of the event under focus is proportional to *Q*^°^(*a, s*)(*N*_*b*|*s*_*/*Σ_*a*∈P_ *N*_*a*|*s*_)*R*^°^(*b* | *a, s*). But the first gene copy could have been sampled in a class *b, s* individual, which occurs with probability *Q*^°^(*b, s*). Then, the probability that the second gene comes from an individual in physiological state *a*, is *N*_*a*|*s*_*/*Σ_*a*∈P_ *N*_*a*|*s*_ and in this case a randomly sampled *a* individual is IBD with the former individual with probability *R*^°^(*a* | *b, s*), whereby the probability of the event under focus is also proportional to *Q*^°^(*b, s*)(*N*_*a*|*s*_*/*Σ_*a*∈P_ *N*_*a*|*s*_)*R*^°^(*a* | *b, s*). Since in both cases we are considering the same event, this yields eq. (49).

Interchanging the dummy variables *a* and *b* in the second summand of eq. (47), using eq. (49), and rearranging yields

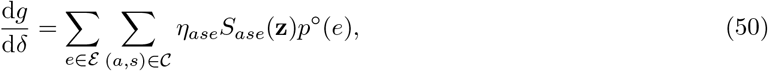

where

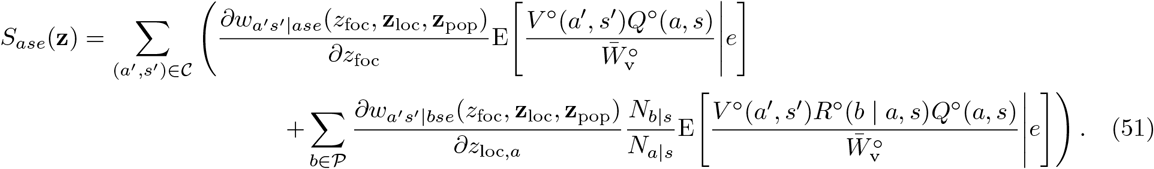

This can be read as an inclusive fitness effect of an individual expressing the mutant deviation *η*_*ase*_ (and evaluated at the resident trait **z**), which aligns with the actor-centered perspective of this concept introduced by Hamilton (1964) (see also Chapter 7 of Rousset, 2004 or Taylor et al., 2007; Scott and Wild, 2023). Specifically, the derivative 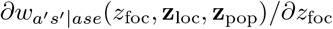 in the first summand (when multiplied by *δη*_*ase*_) represents the additional number of gene copies (or offspring number owing to asexual reproduction) in class (*a*^′^, *s*^′^) produced by a focal actor of class (*a, s*) as a result of it expressing the mutant instead of the resident trait. Meanwhile, the derivative 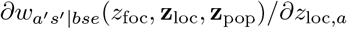 in the second summand can be interpreted as the additional number of class (*a*^′^, *s*^′^) offspring produced by all non-focal local individuals of class (*b, s*) ∈ *C* as a result of the single focal actor of class (*a, s*) expressing the mutant instead of the resident trait. The expectations ensure that each actor is sampled from the stationary distribution of classes in which it resides and that the additional offspring in each class is then weighted with their (stochastic) reproductive value giving their expected asymptotic number of descendants. Relatedness *R*^°^(*b* | *a, s*), meanwhile, guarantees that only non-focal individuals (*b, s*) ∈ *C* sharing a common ancestor with the focal individual (*a, s*) ∈ *C* are included as receptors of the focal trait expression.

From eq. (51) it follows that selection favors an increase in trait component *z*_*ce*_ if

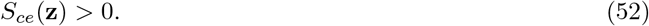

That is, if Hamilton’s marginal rule on trait component *z*_*ce*_ is satisfied. Furthermore, an equilibrium point **z**^*^ of the evolutionary dynamics—a so-called singular point (e.g., Geritz et al., 1997; Rousset, 2004)—must satisfy, for all *ce* ∈ *H*:

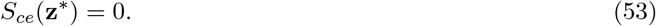

At a singular point, no unilateral deviation in trait value should result in a fitness change and this thus specifies a candidate evolutionary stable trait value. Because *S*_*ce*_(**z**) determines the direction of selection in a resident population, it can also be used to assess the convergence stability of a singular point (Eshel, 1983; Taylor, 1989; Christiansen, 1991; Leimar, 2009) under between-generation environmental fluctuations.

In the absence of between-generation environmental fluctuations, i.e. *w*(*a*^′^, *s*^′^ | *a, s*, **i**, *e*) = *w*(*a*^′^, *s*^′^ | *a, s*, **i**), the derivation of eqs. (50)–(51) from invasion fitness provides an explicit proof of results obtained more heuristically (Ronce et al., 2000, eq. 1), which themselves generalize earlier direct fitness method results (Taylor and Frank, 1996), although in that work the selection gradient was not expressed in terms of an actor-centered inclusive fitness effect (but as in eq. 47 with no environmental fluctuations). Also in the absence of between-generation environmental fluctuations, eqs. (50)–(51) are similar to the inclusive fitness effect obtained for continuous-time processes (Priklopil and Lehmann, 2021, eq. 102). With between-generation environmental fluctuations but in the absence of interactions between relatives, i.e. *R*^°^(*a* | *b, s*) = 0 for all *a, b*, and *s*, eq. (51) can be expressed as

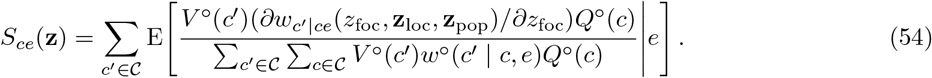

This has the same structure as the classical expression for the derivative of the stochastic growth rate (Tuljapurkar, 1990, eq. 11.2.7), and in particular as eq. (27) of Steinsaltz et al. (2011), up to differences in notation. Consistency with these previous results, obtained by different methods, thus provides a useful check on the present calculations.

The denominator in eq. (54) implies that the contributions of neutral reproductive values and class frequencies cannot, in general, be separated when computing the selection gradient. Such a separation is typically possible in the absence of between-generation environmental fluctuations under models with different levels of complexity (e.g. Taylor, 1990; Taylor and Frank, 1996; Ronce et al., 2000; Rodrigues, 2018b; Priklopil and Lehmann, 2021). The non-separability in eq. (54) holds even though neutral reproductive values and class frequencies are conditionally independent given the current environmental state. The second term in equation (51) makes explicit that the same issue applies to indirect fitness effects. Consequently, the inclusive-fitness effect will generally have to be evaluated using simulations in the presence of between-generation environmental fluctuations, except in special cases permitting further simplification (see section 4.4).

This limitation restricts the range of concrete applications of the inclusive-fitness effect *S*_*ce*_(**z**). Consequently, the main contribution of eq. (51) is interpretative: it shows that, even under different class structures and forms of environmental fluctuation, the direction of a small trait change at any given population state can always be understood in terms of an actor-centered inclusive-fitness effect, consisting of direct and indirect fitness effects resulting from trait change. Thus, regardless of the form of population heterogeneity and underlying stochasticity, the gradual adaptation of quantitative traits can be thought of as resulting from directional kin selection, with the operation of natural selection interpreted in a way closely aligned with that originally proposed by Hamilton (1964). Hence, eq. (51) provides one justification for viewing inclusive-fitness reasoning as an overarching organizing framework for interpreting and formulating hypotheses about adaptation (e.g. Birch, 2017; Levin and Grafen, 2019), one that holds regardless of the complexity of the underlying demographic, ecological, and environmental dynamics.

Readers familiar with perturbation results of the form of eq. (54) for between-generation stochasticity may find it obvious that allowing for interactions between relatives simply adds the second term in eq. (51). Yet this inclusive-fitness effect also accounts for within-generation stochasticity and may appear less obvious from the perspective of the kin-selection literature on selection gradients. In particular, Rousset (2004) derives the selection gradient from the fixation probability of a mutant in a framework that can, in principle, simultaneously incorporate environmental stochasticity, physiological structure, and genetic structure (eq. 11.12 of Rousset, 2004). That result, however, does not coincide with eq. (47), although one might a priori have expected otherwise, since selection gradients obtained from fixation probabilities have been shown to agree with those obtained from perturbations of dominant Lyapunov exponents in a number of cases, and this agreement has been suggested to hold generally (Rousset, 2004, p. 195). The difference can be explained by the fact that computing the derivative of the fixation probability turns out to be proportional to computing the derivative of the growth rate of the average mutant population.^4^ But in a fluctuating environment, the growth rate of the average population differs from the stochastic growth rate (e.g. Tuljapurkar, 1990, p. 47).

This distinction raises the question of which is the relevant measure of Darwinian fitness: fixation probability or invasion fitness? Fully resolving this question is beyond the scope of this paper, but the following observations are in order. If in a finite population mutations to novel trait values are sufficiently rare that fixation or extinction of one mutant lineage typically occurs before another arises, separation-of-timescale results show that a population will spend most of its time fixed for one of a set of possible trait values generated by mutation (Champagnat et al., 2006; Champagnat and Lambert, 2007; Van Cleve, 2015). In this case, evolution can be envisioned as a pure trait-substitution-sequence process, and under mutations with small phenotypic effects, the population will eventually converge to a stochastically stable trait value predicted by a selection gradient computed from the derivative of fixation probability alone (Van Cleve and Lehmann, 2013; Van Cleve, 2015), and this holds for fluctuating environments (Lehmann, 2012). In the regime where the mutation rate is large relative to population size, mutation input becomes comparable to, or exceeds, the inverse duration of selective sweeps, multiple mutant lineages can establish simultaneously, and the isolated-mutant fixation probability no longer provides a sufficient description of evolutionary change. In this regime, not only can morph diversification occur (Wakano and Iwasa, 2013), but the spread of a mutant lineage is described by a branching-process approximation even when the total population is finite (Dawson and Greven, 2014). Thus, directional selection is instead predicted by the derivative of invasion fitness.

### 4.4 Directional derivative for stochastic fitness matrices

#### 4.4.1 Inclusive fitness effect

We now consider a situation where eqs. (50)–(51) can be simplified and made tractable. This is when resident fitness 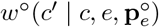 determines a corresponding fitness matrix that is stochastic: 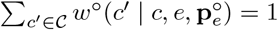 for all *c* and *e*. This can for instance occur when the class structure represents different habitats in the metapopulation like in some classical population genetics models (e.g., Nagylaki, 1992; Karlin, 1982 where *C* = *S* = *ℛ*). In this case under certain demographic conditions, resident fitness 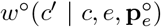 boils down to the probability that an individual reproducing in patch of type *c* produces an offspring migrating to a patch of type *c*^′^ (see eq. 61 for an example). For such a stochastic resident fitness matrix, the unnormalized reproductive values satisfy 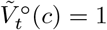 for all *c* ∈ *C*, as can be checked from eq. (25). Therefore:

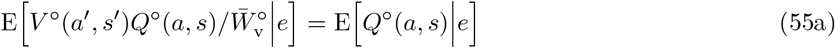

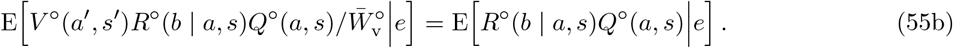

From now on set *q*^°^(*a, s* | *e*) = E [*Q*^°^(*a, s*)|*e*], which is the probability that a randomly sampled individual in a population in state *e* is of class (*a, s*). Meanwhile, E [*R*^°^(*b* | *a, s*)*Q*^°^(*a, s*)|*e*] is the probability that a randomly sampled individual in a population in state *e* is of class (*a, s*) and has a randomly sampled neighbor of class *b* that is IBD to it. Henceforth, we can set *r*^°^(*b* | *a, s, e*) = E [*R*^°^(*b* | *a, s*)*Q*^°^(*a, s*)|*e*] */q*^°^(*a, s* | *e*), which is the probability that in a population in environmental state *e* and given an individual has been sampled in class (*a, s*), a randomly sampled neighbor of class *b* is IBD to the former individual. By taking expectation over eq. (49) we have the following relationship:

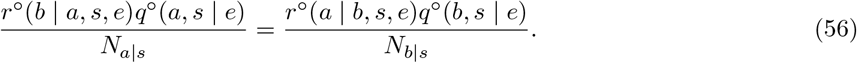

When eq. (55) holds, then by using eq. (56) we can simplify eq. (51) to

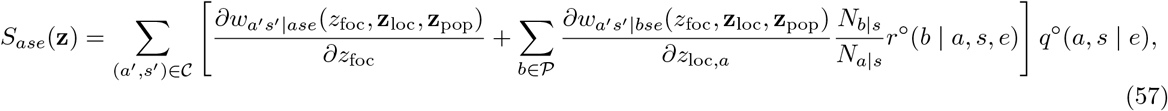

where the class frequencies satisfy the recursion

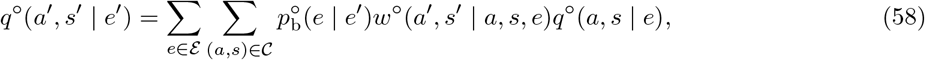

while the relatedness coefficients satisfy

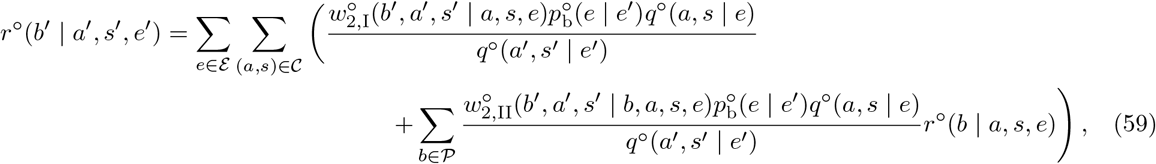

with 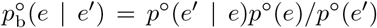 being the probability that a population in state *e*^′^ was in state *e* in the previous time step (Appendix B.1 for a proof of these two recursions). The first ratio in parenthesis can be interpreted as the probability that two offspring individuals, one of physiological state *a*^′^ the other of state *b*^′^, both sampled when the group and environmental states are (*s*^′^, *e*^′^), descend from a single individual in class *a* that reproduced in environmental state (*s, e*). In this case, the IBD probability between the two offspring is one. The second ratio in parenthesis can be interpreted as the probability that two individuals, one of physiological state *a*^′^ the other of state *b*^′^, both sampled in environmental state (*s*^′^, *e*^′^) both descend from two distinct individuals in environmental state (*s, e*), one in class *a* the other in class *b*. In this case, the IBD probability between the two offspring is *r*^°^(*b* | *a, s, e*).

Eqs. (57)–(59) show that, for a stochastic fitness matrix, the selection gradient can be simplified and evaluated by solving a linear system of equations. Thus, directional selection can now be ascertained without simulating the lineage process. Additionally, 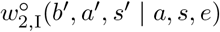 and 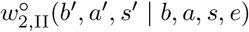 can usually be evaluated straightforwardly using identity-by-descent arguments (e.g., Ronce et al., 2000 and Ohtsuki et al., 2020). Note that a special case in which eq. (57) can be systematically applied is when there is no class structure at all (in which case eq. (58) is not needed and eq. (59) can be markedly simplified). This case covers classical models of the evolution of sex ratio, dispersal, or altruism in spatially structured populations (e.g., Hamilton and May, 1977; Frank, 1998; Taylor and Irwin, 2000; Rousset, 2004), which can thus all be analyzed under conditions of between-generation environmental fluctuations.

#### 4.4.2 Application: lottery model with spatial heterogeneity under soft selection

We now provide an application of eqs. (57)–(59) by extending the ecological scenario of Svardal et al. (2015) to allow for interaction between relatives. This scenario considers an island model with spatial heterogeneity where *S* = *ℛ* = {1, 2, …, *H*} represents a finite number *H* of habitat types that patches can occupy, and where the frequency of habitat of type *s* is *p*^°^(*s* | *e*) = *π*(*s*) and is fixed. We assume that each patch is of equal and finite size *N*, instead of being of infinite size as in the original scenario of Svardal et al. (2015), but otherwise follow their life-cycle, which is as follows. (1) Each adult in a patch produces a very large offspring number *f* (*z, s, e*) that depends on its trait *z*, the patch state *s* in which the individual resides and the current population state *e*. (2) After reproduction each adult individual survives with probability *γ* or dies with complementary probability. (3) Regulation occurs among the local offspring pool and brings back the local number of offspring proportionally to the vacated breeding spots, *N* (1 − *γ*) on average (“soft selection”). (4) Each of these offspring either disperses with probability *d* or stays local and then competes to be recruited.

It follows from these assumptions and the fact that there is no physiological state (and thus *C* = ℝ) that individual fitness can be written as

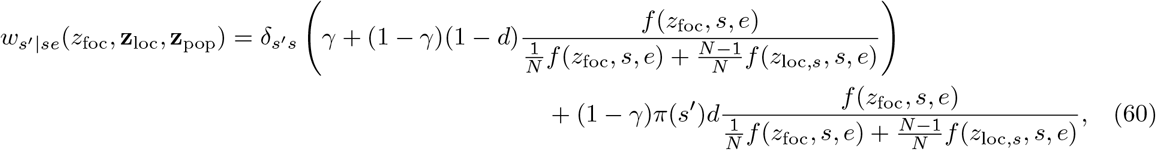

where 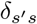 is the Kronecker-delta. When *N* → ∞, this is equivalent to the fitness determined by eq. (4a)–(4b) of Svardal et al. (2015). Neutral fitness 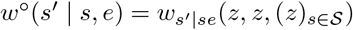 reduces to

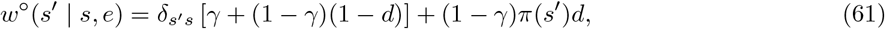

which specifies a stochastic fitness matrix:

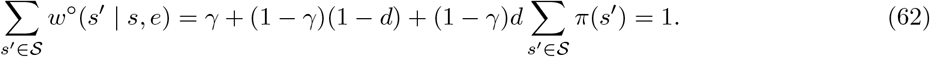

Thus, we can apply eq. (57) to the present scenario, which, in the absence of environmental heterogeneity is a special case of the soft selection model of Ohtsuki et al. (2020, section 4 and eq. F.23). Their results also imply that we can write

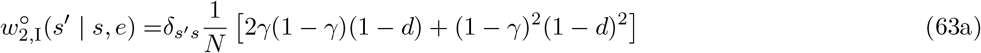

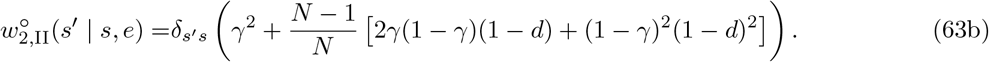

We now have all the ingredients to compute the selection gradient for the present scenario, which reduces to

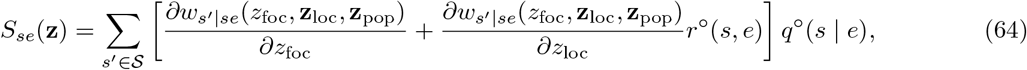

where, owing to eqs. (60)–(63), the direct fitness effect is

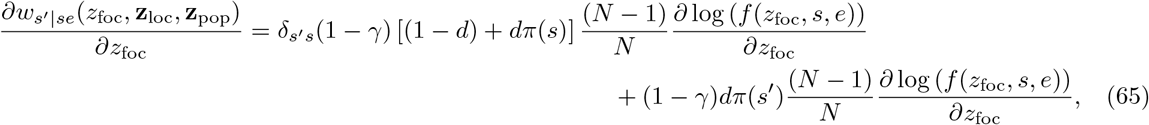

the indirect fitness effect is

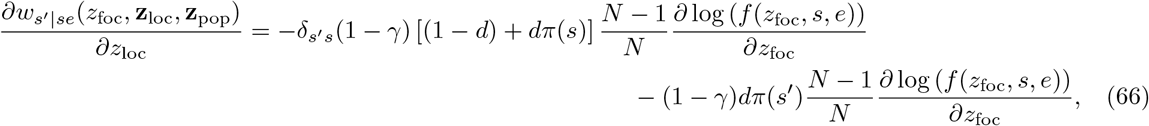

and

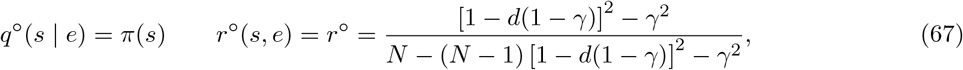

which are both shown in Appendix B.2. The constant relatedness obtained here corresponds to that derived earlier for survival in the island model in the absence of environmental fluctuations (e.g. Pen, 2000; Taylor and Irwin, 2000).

Substituting eqs. (65)–(67) into eq. (64) and in turn into eq. (50) by setting *η*_*ce*_ = 1 yields

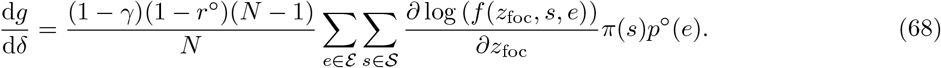

This result is fully consistent with eq. (11) of Svardal et al. (2015) and shows that the condition determining the location of the singular point, i.e., the value of *z*^*^ satisfying d*g/* d*δ* = 0, is unaffected by the finiteness of patches and the occurrence of kin selection. The reason is that fecundity *f* (*z*_foc_, *s, e*) depends only on the actor’s trait, not on the traits of other individuals in its patch, and so the indirect effect, eq. (66), is similar to, but opposite in sign to, the direct effect, eq. (65). Eq. (68) will no longer hold if some of the assumptions underlying the scenario are relaxed, for instance if survival *γ*(*e*) is allowed to depend on the environmental state or if fecundity depends on the traits of other patch members.

## 5 Discussion

The quantitative analysis of natural selection begins with a measure of Darwinian fitness. Since at least Fisher (1930), it has been recognized that, in large, ideally infinite, populations, Darwinian fitness is determined by the geometric growth rate of a mutant in a resident population. In this perspective, Darwinian fitness is identified with invasion fitness and determined by an appropriate stochastic growth rate in a fluctuating environment.

This paper builds on standard results from multitype branching processes in random environments and applies them to group-structured populations subject to multiple sources of discrete heterogeneity: heterogeneity both within and among groups, as well as stochasticity both within and between generations. Invasion fitness is then determined by a stochastic growth rate governed by products of matrices describing expected reproductive output within generations (eq. 9). This yields two biological representations of invasion fitness. The first expresses it as the long-term geometric mean of the expected individual fitness of a mutant carrier, where the average is taken over all demographic, genetic, physiological, and environmental heterogeneities and stochasticities within generations (eq. 21). The second expresses it as the long-term geometric mean of reproductive value–weighted average individual fitness across these same contexts, effectively filtering out non-heritable, generation-specific contributions to fitness (eq. 32). Despite their different constructions, both representations of invasion fitness support a common genealogical perspective: invasion fitness corresponds to a geometric mean across generations of the expected fitness of a randomly sampled lineage member within each generation. In this sense, the “gene lineage is the bearer of fitness” (Akçay and Van Cleve, 2016), as has long been recognized from the gene-centered perspective on evolution (Hamilton, 1963; Dawkins, 1978; Haig, 2012). Moreover, because lineage members are identical-by-descent, natural selection arising through differential lineage fitness in the presence of interactions among lineage members cannot be divorced from kin selection, any more than natural selection can be separated from adaptation.

A useful feature of the reproductive value–weighted representation of invasion fitness is that it enables a direct computation of the phenotypic derivative of the stochastic growth rate when traits can be expressed in a context specific way and interactions occur between relatives. The resulting directional derivative can be written entirely in terms of resident population quantities; namely, individual fitness differentials, class densities, reproductive values, and relatedness (eq. 51). This result extends a classical expression for the derivatives of the stochastic growth rate (Tuljapurkar, 1990, eq. 11.2.7, Steinsaltz et al., 2011, eq. 27) by explicitly incorporating interactions among relatives arising from limited genetic mixing and small group size thus including demographic stochasticity. The resulting selection gradient not only decomposes into direct and indirect fitness effects, but yields a genuine actor-centered representation of directional selection. Overall, this thus provides a bridge between invasion processes in stochastically fluctuating environments and inclusive fitness theory (e.g. Hamilton, 1964; Frank, 1998; Rousset, 2004).

A difficulty arises, however, in the presence of between generation stochasticity. Unlike under other forms of heterogeneity, class densities, reproductive values, and relatedness are themselves random variables and thus stochastic. As a consequence of selection fluctuating across generations, they cannot be decoupled and solved independently via systems of linear equations, as is possible in the deterministic case under within generation heterogeneity of arbitrary complexity (Rousset, 2004; Priklopil and Lehmann, 2021). This interdependence limits analytical tractability: the selection gradient typically cannot be obtained in closed form and “even with a formula in hand we must compute derivatives of the stochastic growth rate by numerical simulation, which is subject to both sampling error and bias” (Steinsaltz et al., 2011). This latter work provides bounds on such errors that can be used in applications. Thus, while the derived expression for the directional phenotypic derivative generalizes marginal Hamilton’s rule to stochastic environments with class structure– and is therefore useful for understanding the nature of directional natural selection on quantitative traits from a biological perspective–its practical applicability is constrained. Nevertheless, there is a special case in which tractability is regained. This occurs when the expected per-generation fitness matrix is stochastic; in that case, the selection gradient simplifies and takes a form conceptually similar to that in the absence of stochastic fluctuations (eq. 57). Although this situation is not generic, it is biologically relevant. For instance, a number of metapopulation scenarios with heterogeneous habitats fall into this category (section 4.4.2 for an example).

Despite addressing a range of biological scenarios, the present analysis has several limitations and poses some challenges for future work. First, the model assumes that fitness depends only on the global environmental state of the parental generation and not on that of the offspring generation. This assumption was adopted for simplicity of notation and would not be difficult to relax. Second, the model assumes haploid reproduction. Diploidy and sexual reproduction can be incorporated by extending the class structure and, under additive gene action, primarily affect directional selection through a rescaling of fitness differentials and an appropriate reinterpretation of relatedness (Rousset, 2004). Nevertheless, it would be useful to work out these extensions explicitly. Third, the model is restricted to a discrete set of classes. Extending the framework to continuous class structures and continuous-time stochastic demographic processes would be a natural and valuable generalization, particularly for studying life-history evolution. Fourth, the analysis is limited to first-order derivatives of the stochastic growth rate. Access to second-order derivatives would allow a full classification of adaptive-dynamics singularities (e.g. Geritz et al., 1997; Brännström et al., 2013; Avila and Mullon, 2023). This could be achieved by extending the analysis of Ohtsuki et al. (2020) to the present setting. Fifth, the model assumes an infinite resident population size, and the connection with the derivative of fixation probability in finite populations should be developed to determine the conditions on mutation rates and population size under which gradual evolution in finite populations is governed by the derivative of invasion fitness rather than by the derivative of fixation probability (see also the last paragraph of section 4.3). Finally, approximations could be developed to simplify the general expression for the selection gradient and make it more tractable—for instance, by applying a weak-noise approximation (Tuljapurkar, 1990) or by adopting semi-deterministic local demographic dynamics (Mullon and Lehmann, 2018). We thus hope that the present analysis will motivate further theoretical work on evolution in populations experiencing both limited dispersal and environmental fluctuations across generations.

## Acknowledgments

I am very grateful to David Steinsaltz for setting me straight by explaining that the unnormalized neutral reproductive value dynamic is a null-recurrent Markov chain and thus has no stationary distribution. I am also very grateful to Jeremy Van Cleve for useful discussions and comments on earlier versions of the manuscript. Finally, many thanks to Ludovic Maisonneuve for spotting oddities.

## Appendix A Neutral recursions

### Appendix A.1 Class frequencies

We here derive eq. (45). To get there, denote by 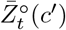 the expected number of individuals at time *t* in class *c*^′^ ∈ *C* that descend from a single resident ancestor individual, tagged at time *t* = 0 as the progenitor of a resident lineage. This random variable satisfies the recursion

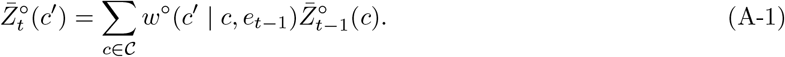

The frequency of class *c*^′^ individuals of the resident lineage at time *t* is 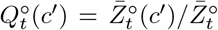, where 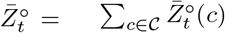, and using this along eq. (A-1) produces

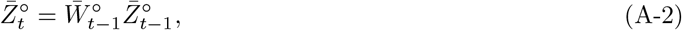

where

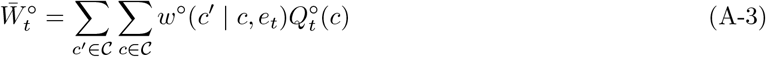

is mean neutral fitness (the latter two equations are same as eqs. (16)–(17) under neutrality and noting that 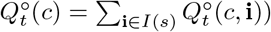. Now from the definition of class frequencies and eq. (A-2) we finally get

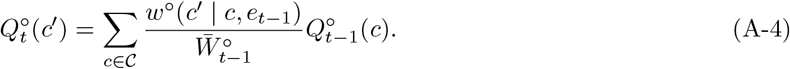

### Appendix A.2 Relatedness

We now derive eq. (46). For this, let us recall the definition of relatedness, eq. (43), and first write it regardless of the strength of selection on the mutant type, as

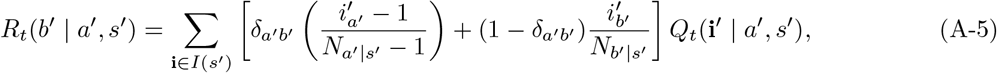

where *δ*_*ij*_ denotes the Kronecker delta and owing to eq. (18) we have

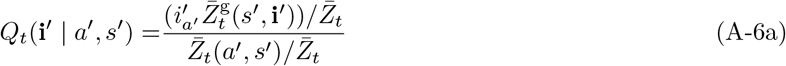

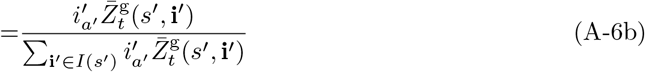

with

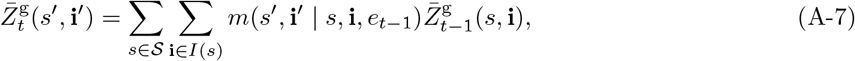

which follows from eq. (13).

Building on Ohtsuki et al. (2020, eq. 11), let us now generalize eq. (15) to the second moment of fitness, and observe that

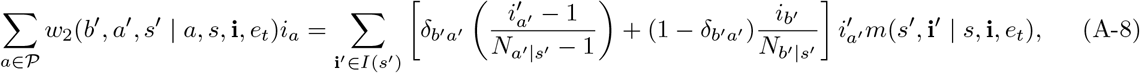

where *w*_2_(*b*^′^, *a*^′^, *s*^′^ | *a, s*, **i**, *e*_*t*_) is the expected number of offspring of class (*a*^′^, *s*^′^) produced by a single mutant parent reproducing in class (*a, s*) in a group in genetic state **i** in a population in environmental state *e*_*t*_, and where the offspring have a randomly sampled neighbor in physiological state *b*^′^ that is also a mutant. When *b*^′^ = *a*^′^, the neighbour must be a distinct individual from the offspring of the focal, and if there are 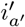 mutants in class *a*^′^ in a group in state *s*^′^, a random neighbour is a mutant with probability 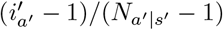.

On substituting eqs. (A-6)–(A-7) into eq. (A-5) and using eq. (15) and eq. (A-8), we get that

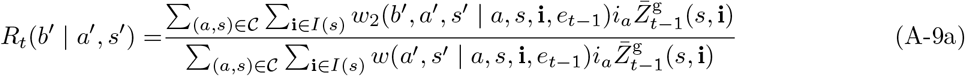

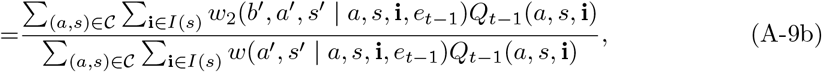

where eq. (18) was used in the second line. In the absence of physiological class structure and environmental heterogeneity, this is similar to Ohtsuki et al. (2020, eq. C.11).

Let us again build on Ohtsuki et al. (2020, eq. 28) and observe that under neutrality we can write

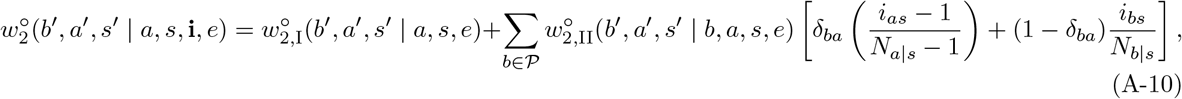

where *w*^°^ (*b*^′^, *a*^′^, *s*^′^ | *a, s, e*) is the expected number of offspring in class (*a*^′^, *s*^′^) that descend from a parent reproducing in class (*a, s*) and that have a random neighbor in physiological state *b*^′^ that also descends from that parent (where all this happens in the neutral process). Namely, this fitness component measures the number of sibling pairs in physiological state *a*^′^ and *b*^′^ produced by a focal parent. Fitness component 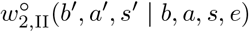 is the expected number of offspring in class (*a*^′^, *s*^′^) that descend from a focal parent reproducing in class (*a, s*) and that have a random neighbor in physiological state *b*^′^ that descends from any given neighbour in physiological state *b* of the focal parent.^5^

The above fitness moments are perhaps better understood biologically by considering the similarity and difference to individual fitness *w*^°^(*a*^′^, *s*^′^ | *a, s, e*) and noting that:

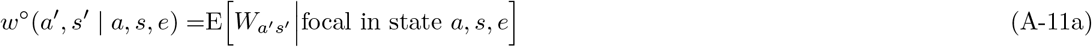

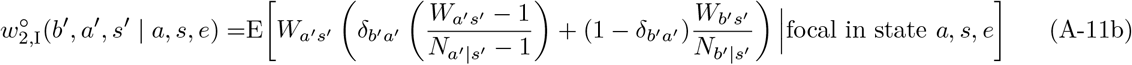

and

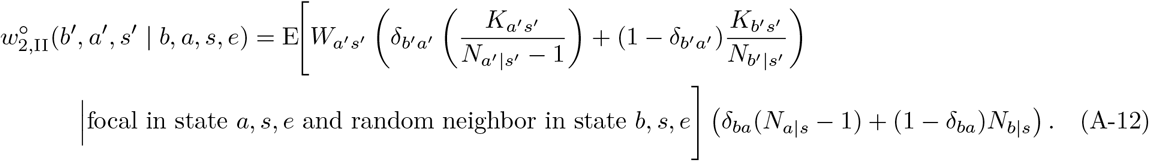

Here, *W*_*as*_ denotes the random number of successful offspring in physiological state *a* in a group in state *s* produced by a focal individual (whose state is characterized by the conditioning in the expectation), and *K*_*as*_ denotes the random number of successful offspring in physiological state *a* in the same group in state *s* produced by a random neighbor of the focal individual. The conditional expectations are taken over all within-generation stochasticity affecting the fitness of interacting individuals [for an explicit example of computations of such fitness moments, see Ohtsuki et al., 2020, Appendix F.2]. Eq. (A-11) makes explicit that within-generation stochasticity is fully accounted for in the construction of the individual fitness components of the model. In turn, these expectations may depend on the means, variances, and higher moments of the fecundity distributions of interacting individuals (Lehmann and Balloux, 2007, eq. A.6), as is the case under within-generation bet-hedging (Gillespie, 1974, 1977).

On substituting eq. (24) and eq. (A-10) into the second line of (A-9), evaluating under neutrality and simplifying, relatedness is seen to satisfy the recursion

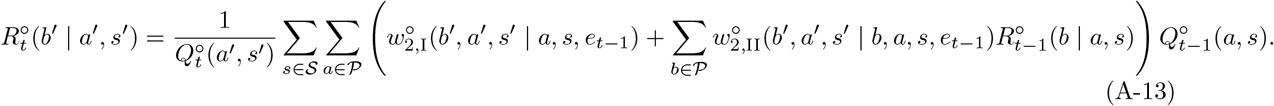

## Appendix B Neutral stochastic fitness matrix

### Appendix B.1 Recursions

#### Appendix B.1.1 Class frequencies

We here derive eq. (58). First note that because the fitness matrix is assumed to be stochastic, we have from eq. (A-3) that 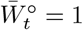 for all *t* 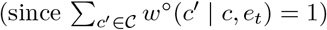, whereby eq. (A-4) yields

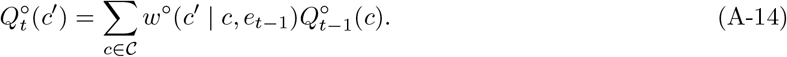

Now, we first observe that using the law of total conditional expectation, and conditioning additionally on the environmental state (*e*_*t*−1_ = *e*) in the parental generation, we can write

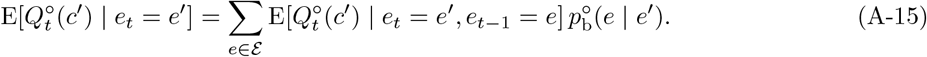

Using eq. (A-14), we have

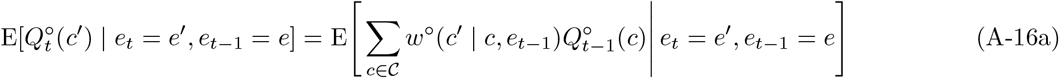

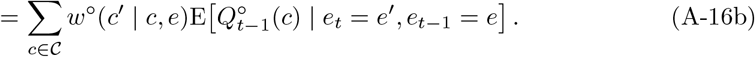

Since the environmental process is exogenous and Markovian, 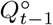 and *e*_*t*_ are conditionally independent given *e*_*t*−1_. Therefore,

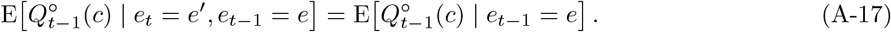

Setting E [*Q*^°^(*c*)|*e*] = *q*^°^(*c* | *e*) and using eqs. (A-15)–(A-17) at stationarity, we then obtain

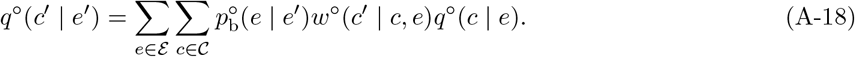

#### Appendix B.1.2 Relatedness

We now derive eq. (59) using the same argument as in the previous section. First, by conditioning on the environmental state in the parental generation, we have

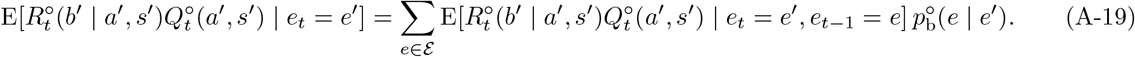

Using eq. (A-13), we obtain

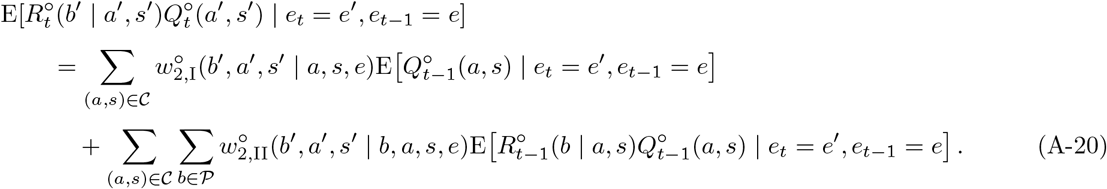

Since the environmental process is exogenous and Markovian, 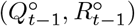 and *e*_*t*_ are conditionally independent given *e*_*t*−1_, and thus we have

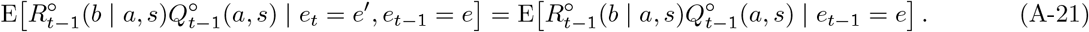

At stationarity, set E [*Q*^°^(*c*)|*e*] = *q*^°^(*c* | *e*) and *r*^°^(*b* | *a, s, e*) = E [*R*^°^(*b* | *a, s*)*Q*^°^(*a, s*)|*e*] */q*^°^(*a, s* | *e*) and using eq. (A-17) and eqs. (A-19)–(A-21), we obtain

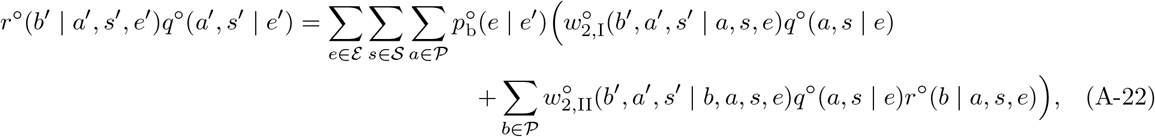

which can be expressed as eq. (59).

### Appendix B.2 Lottery model with spatial heterogeneity under soft selection

We here derive eq. (67). By substituting eq. (61) into eq. (58), we get

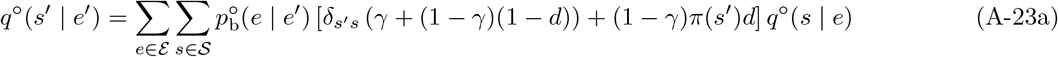

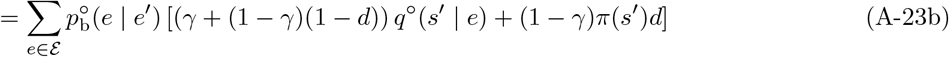

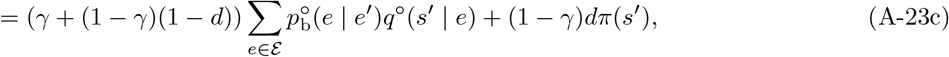

where in the second equality we used Σ_*s∈S*_ *q*^°^(*s* | *e*) = 1, and in the third 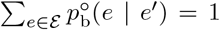. Now by substituting

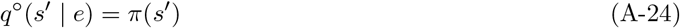

into eq. (A-23c) it is clear that this is a solution to the recursion for all *e* ∈ *E* and *s*^′^ ∈ *S*. Moreover, this is the unique solution when (1 − *γ*)*d >* 0, as can be seen by writing eq. (A-23c) in vector form as

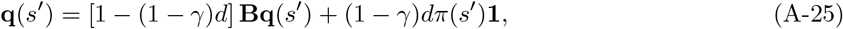

where **q**(*s*^′^) = {*q*^°^(*s*^′^ | *e*)}_*e*∈*E*_, **1** is a vector of 1 of the same dimension, and **B** is a row stochastic matrix (**B1** = **1**) with elements 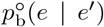. Since (**I** − [1 − (1 − *γ*)*d*] **B**) **1** = (1 − *γ*)*d***1**, we have (1*/*(1 − *γ*)*d*)**1** = (**I** − [1 − (1 − *γ*)*d*] **B**)^−1^ **1** and it follows that **q**(*s*^′^) = *π*(*s*^′^)**1** (where **I** is the identity matrix).

Let us now turn to the relatedness recursion, where eq. (59) under the present scenario reduces to

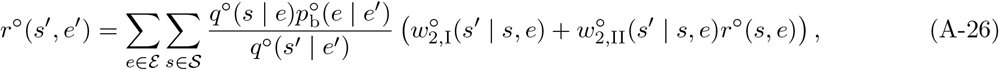

and on substituting eq. (63) yields

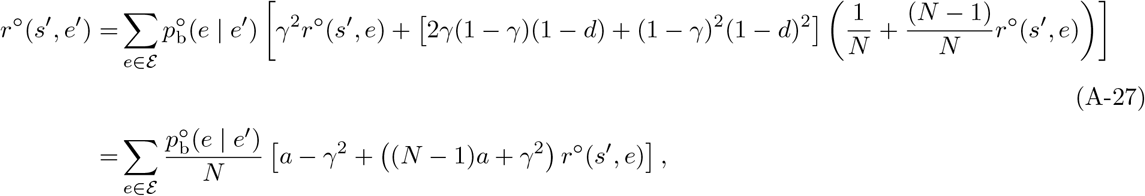

where *a* = (1 − *d*(1 − *γ*))^2^. Expressing this recursion in vector form by using **r**(*s*^′^) = {*r*^°^(*s*^′^, *e*)}_*e*∈*E*_, we have

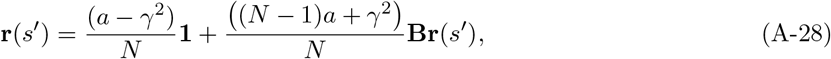

and since (**I** – [((*N* − 1)*a* + *γ*^2^) */N*] **B**) **1** = (1 – ((*N* − 1)*a* + *γ*^2^) */N*) **1**, it follows that **r**(*s*^′^) = *r*^°^**1**, where

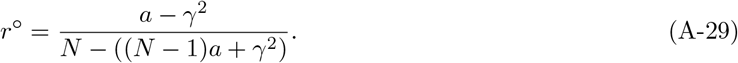

#### Box 1.

**Individual fitness**

To illustrate the concept of individual fitness *w*^°^(*c*^′^ | *c, e*), consider the model of an age-structured population under limited dispersal with exactly one individual per group, as described by Ronce and Promislow (2010), but with environmental fluctuations added. In this model, newborns can migrate between groups, but adults remain sessile. The physiological state space *P* = {1, 2, …, *T* } then represents the set of ages an individual can occupy and *T* is the maximum lifespan. Since there are no local fluctuations in group size, the group state space corresponds directly to the physiological state space, which in turn determines class space: *C* = *S* = *P*. Let *f* (*a, e*) denote the fecundity of an individual of age *a* in environmental state *e* and *γ*(*a, e*) the corresponding survival probability. Finally, let *d* be the migration rate of juveniles out of their natal patch. For this model, the individual fitness through survival by an individual of age *a* in a population in environment *e* is for *a < T*

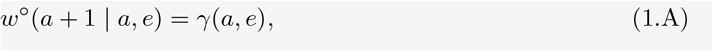

and otherwise *w*^°^(*a*^′^ | *a, e*) = 0 for *a*^′^ ∈/{1, *a* + 1}. Meanwhile, the production of age class one individuals by an individual of age *a* in *e* is

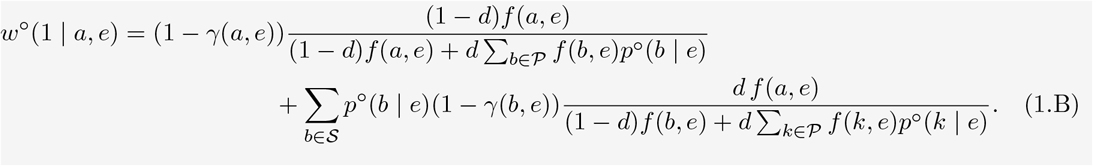

Here, *p*^°^(*b* | *e*) is the frequency of groups in state *b* in the metapopulation when the environment is *e*, and which determine the profile 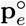. This distribution can be obtained from eq. (3)–(4) with transition probability specified by eq. (2), itself constructed by specifying an environmental process (eq. (1)), and noting that *γ*(*a, e*) = *p*^°^(*a* + 1 | *a, e*) is the probability that a group with an individual of age *a* turns into a group with an individual of age *a* + 1 in state *e*, while 1 − *γ*(*a, e*) = *p*^°^(1 | *a, e*) is the probability that a group with an individual of age *a* turns into a group with a newborn. In the absence of global environmental heterogeneity, eq. (1.B) corresponds to eqs. (2.2)–(2.3) in Ronce and Promislow (2010).

## Footnotes

1 When there is exactly one mutant individual of physiological state *a* in group state *s*, the state (*s*, i) of a group is represented by (*s*, u_*a*_), where u_*a*_ is the unit vector with a 1 in position *a* and zeros otherwise. The set of all such possible single-mutant states is in one-to-one correspondence with the class space defined as C = {(*a, s*) ∈ P × S | *N*_*a*|*s*_ ≥ 1}, since u_*a*_ ∈ *I*(*s*) if and only if *N*_*a*|*s*_ ≥ 1, and each state (*s*, u_*a*_) therefore uniquely identifies the pair (*a, s*).

2 The equivalence can be shown as follows. Suppose, Pr[*ϕ*_*c*_(e) = 1] = 1, then the only value *ϕ*_*c*_(e) can take under the probability distribution is one, and thus E[*ϕ*_*c*_(e)] = 1, i.e. Pr[*ϕ*_*c*_(e) = 1] = 1 =⇒ E[*ϕ*_*c*_(e)] = 1. Consider now the other direction and let 1 − *ϕ*_*c*_(e) stand for the probability of non-extinction on realization *e*. By eq. (1.16) of Karlin and Taylor (1975), Pr[1 − *ϕ*_*c*_(e) *>* 0] ≤ E[1 − *ϕ*_*c*_(e)]*/ϵ* for all *ϵ >* 0. If E[*ϕ*_*c*_(e)] = 1 holds, then E[1 − *ϕ*_*c*_(e)] = 0, hence Pr[1 − *ϕ*_*c*_(e) *> ϵ*] = 0 for any *ϵ >* 0. So Pr[1 − *ϕ*_*c*_(e) *>* 0] = 0 almost surely, whereby Pr[*ϕ*_*c*_(e) = 1] = 1, i.e. E[*ϕ*_*c*_(e)] = 1 =⇒ Pr[*ϕ*_*c*_(e) = 1] = 1.

3 While the demographic model of Steinsaltz et al. (2011) is not framed in terms of limited dispersal with genetic states and ignores within-generation demographic stochasticity, their mathematical machinery can nevertheless be applied to the present model, as it applies to any model of the form given by eqs. (10)–(13), i.e. the growth of a population structured into a finite number of states under a Markovian stochastic environment that itself has a finite number of states. This application is possible thanks to the remarkable branching-process result encapsulated by eq. (9), which provides a bridge between within- and between-generation stochasticity.

4 The growth rate of the average population can be obtained by computing the leading eigenvalue of the megamatrix (Tuljapurkar et al., 2003), which, in the present context, is a matrix whose elements are the process matrices M(*e*)–themselves consisting of elements *m*(*s*^*′*^, i^*′*^ | *s*, i, *e*)–weighted by the transition probability *p*^*°*^(*e*^*′*^ | *e*). Calculations then show that the derivative of the megamatrix eigenvalue can be expressed in a form analogous to eq. (11.12) of Rousset (2004) in the context of the island model.

5 In Ohtsuki et al. (2020, eq. 28), the term corresponding to the second component in eq. (A-10) does not involve any denominator with number of individuals. This is because they express the decomposition in eq. (A-10) in terms of the expected number of offspring in class (*a*^*′*^, *s*^*′*^) that descend from a parent reproducing in class (*a, s*) and that have a random neighbor in physiological state *b*^*′*^ that descend from a randomly sampled parent (instead of any parent) in physiological state *b*. The present formulation leads to more compact expressions for relatedness when the population is class structured, hence the different formalization.

